# The BLIMP1 – EZH2 nexus in a non-Hodgkin lymphoma

**DOI:** 10.1101/606749

**Authors:** Kimberley Jade Anderson, Árný Björg Ósvaldsdóttir, Birgit Atzinger, Gunnhildur Ásta Traustadóttir, Kirstine Nolling Jensen, Aðalheiður Elín Lárusdóttir, Jón Þór Bergþorsson, Ingibjörg Harðardóttir, Erna Magnúsdóttir

## Abstract

Waldenström’s macroglobulinemia (WM) is a non-Hodgkin lymphoma, resulting in antibody-secreting lymphoplasmacytic cells in the bone marrow and pathologies resulting from high levels of monoclonal immunoglobulin M (IgM) in the blood. Despite the key role for BLIMP1 in plasma cell maturation and antibody secretion, its potential role in WM cell biology has not yet been explored. Here we provide evidence of a crucial role for BLIMP1 in the survival of WM cells and further demonstrate that BLIMP1 is necessary for the expression of the histone methyltransferase EZH2 in both WM and multiple myeloma. The effect of BLIMP1 on EZH2 levels is post translational, at least partially through the regulation of proteasomal targeting of EZH2. Chromatin immunoprecipitation analysis and transcriptome profiling suggest that the two factors co-operate in regulating genes involved in cancer cell immune evasion. Co-cultures of natural killer cells and WM cells further reveal that both factors participate directly in immune evasion, promoting escape from natural killer cell mediated cytotoxicity. Together, the interplay of BLIMP1 and EZH2 plays a vital role in promoting the survival of WM cells.

## Introduction

Waldenström’s macroglobulinemia (WM) is a rare plasma cell dyscrasia with around 3-6 people per million diagnosed annually world wide [1–4]. It is characterised by the expansion of a monoclonal population of malignant cells in the bone marrow with a lymphoplasmacytic character, that is, cellular phenotypes ranging from that of B-lymphocytes to overt plasma cells that exhibit hypersecretion of immunoglobulin M (IgM) [5]. A large proportion of WM symptoms arise because of high levels of IgM paraprotein in patients’ blood and tissues [6]. Curiously, over 90% of WM tumours carry an activating mutation in the signaling adaptor MYD88, typically L265P that serves as a key oncogenic driver in the disease [7–9].

The transcription factor B-lymphocyte induced maturation protein-1 (BLIMP1) drives plasma cell differentiation, mediating transcriptional changes via the recruitment of co-factors to chromatin [10–12]. During plasma cell maturation, BLIMP1 represses key B-lymphocyte identity factors and signalling mediators [13, 14], while simultaneously driving plasma cell specific gene expression and antibody secretion [15–17]. Depending on the mouse model system used, BLIMP1 appears to be required for the survival of long-lived plasma cells in the bone marrow and promotes multiple myeloma (MM) cell survival [16, 18–20]. Conversely, BLIMP1 functions as a tumour suppressor in diffuse large B cell lymphoma (DLBCL), consistent with its repression of proliferation genes during plasma cell differentiation [15, 21–23]. Consistent with a potential tumour suppressor role, WM tumours harbour frequent heterozygous losses of *PRDM1*, the gene encoding BLIMP1 [24]. However, BLIMP1 is expressed in a subset of WM lymphoplasmacytic cells [25, 26], in line with its necessity for antibody secretion [17, 19, 27], a critical aspect of WM pathology. Furthermore, BLIMP1 is induced downstream of toll like receptor engagement and MYD88 [28, 29] and *PRDM1* mRNA is elevated in tumours harbouring the MYD88^L265P^ mutation [30], which is associated with poorer prognosis in WM [31].

Also important in plasma cell differentiation, enhancer of zeste 2 (EZH2) is both a physical and genetic interaction partner of BLIMP1 [17, 32]. The interaction was first suggested in mouse primordial germ cells, where BLIMP1 and EZH2 share a highly overlapping set of binding sites [33, 34]. EZH2 is the catalytic component of the polycomb repressive complex 2, placing methyl groups on lysine 27 of histone 3, typically tri-methylation (H3K27me3), to repress transcription [35, 36]. While EZH2 is essential for embryonic development [37], it has frequent activating mutations in DLBCL, and is commonly overexpressed in MM, making it a promising therapeutic target [38–40]. Surprisingly, while aberrant regulation of histone modifications has been implicated in WM pathogenesis [41], the role of EZH2 is yet to be investigated.

In this study, we examined the role of BLIMP1 in WM, and its potential interplay with EZH2. We demonstrate for the first time that BLIMP1 regulates WM cell survival and maintains EZH2 protein levels. We identify a large overlap in transcriptional targets of the two factors and show that they repress transcription on an overlapping set of genes in a parallel fashion. In a highly novel finding, we reveal roles for BLIMP1 and EZH2 in evasion from natural-killer (NK) cell mediated cytotoxicity, with BLIMP1 suppressing NK cell activation in response to WM cells and both factors suppressing NK cell mediated WM cell death. Thus, inhibition of BLIMP1 or EZH2 could be a promising therapeutic strategy. Together, our data highlights the multifaceted roles of the BLIMP1-EZH2 nexus in WM cell survival.

## Materials and Methods

### RNA isolation, cDNA synthesis and RT-qPCR

Cells were lysed in TRIsure reagent (#BIO-38032, Bioline, London, UK), and RNA was extracted according to the manufacturer’s instructions. cDNA synthesis and RT-qPCR are described in supplementary materials and methods.

### Viability and reduction assays

The percentage of live cells was determined by trypan blue exclusion assay (Thermo Fisher Scientific, USA). The per cent reduction was determined by resazurin assay (#sc-206037, Santa Cruz Biotechnology, CA, USA).

### Proteasomal inhibition

To inhibit proteasome activity, the cells were treated for 4 h with 5µM MG-132 (#sc-201270, Santa Cruz Biotechnology, dissolved in DMSO), 20 h post dox addition.

### RNAseq and ChIPseq

Prior to RNAseq, The *PmiR2* and *NTmiR* RP cells were treated with 0.2µg/mL dox for 48 h. RP cells were treated with 300nM tazemetostat (EPZ-6438, #S7128, Selleckchem, Munich, Germany) or DMSO for 48 h. Transcription factor ChIP for BLIMP1 and EZH2 was performed as previously described [33, 42, 43]. Histone ChIP for pan-H3 and the H3K27me3 mark was performed as previously described [44].

### NK cell isolation, degranulation and cytotoxixity assays

NK cells were isolated from heparinised buffy coats obtained from healthy human donors, who all provided informed consent, provided by the Icelandic Blood Bank (ethical approval #06-068).

Additional details for Western blotting, immunofluorescence, reduction assays, RNAseq, ChIPseq, NK cell isolation, degranulation and cytotoxicity assays are described in Supplementary Materials and Methods.

## Results

### BLIMP1 is important for cell survival in Waldenström’s macroglobulinemia

As BLIMP1 is expressed in a subset of WM patients’ lymphoplasmacytic cells [25, 26], and given its crucial roles in antibody secretion and plasma cell differentiation, we wanted to determine whether it plays a role in WM cell biology. We compared BLIMP1 expression in the myeloma cell line OPM-2 [45] to that of three WM cell lines RPCI-WM1 (RP), MWCL-1 (MW) and BCWM.1 (BC) by immunofluorescence staining (Fig. 1A). All of the RP cells expressed BLIMP1, while the MW and BC cells had more heterogeneous expression, with 43% and 18% of the cells expressing high levels of BLIMP1 respectively (Fig. 1A right panel). This heterogeneity of BLIMP1 expression is interesting because the antibody-secreting cells responsible for the IgM associated pathology of WM are likely dependent on BLIMP1 for secretion. We subsequently chose to focus mainly on the RP cell line as it had the most uniform BLIMP1 expression.

**Figure 1:**
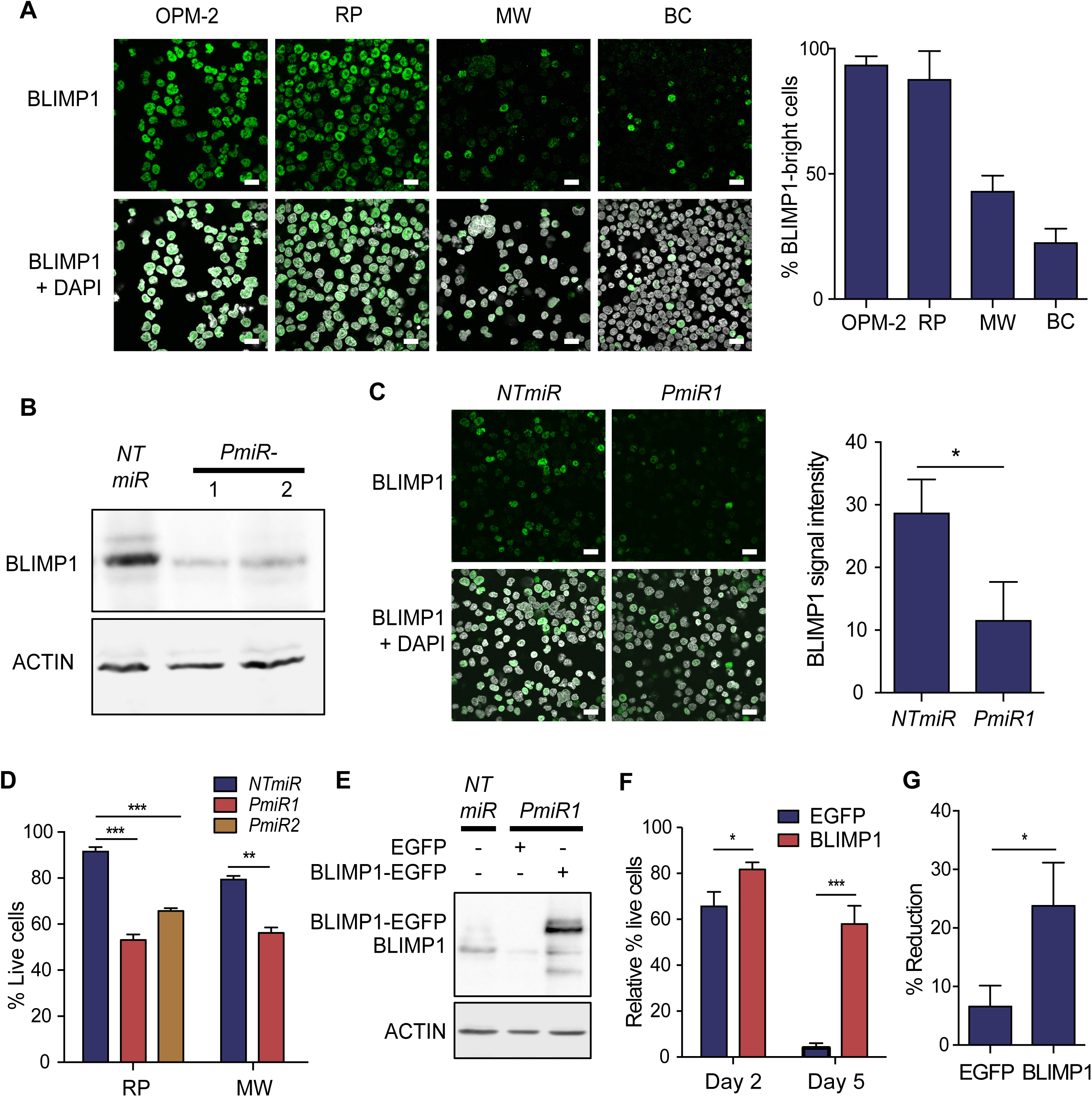
BLIMP1 promotes the survival of WM cells. **(A)** BLIMP1 expression in the myeloma cell line OPM-2 and in WM cell lines RP, MW and BC as detected by immunofluorescence staining, with a bar graph representing percentage of BLIMP1^bright^ cells. Scale bars represent 20µm. **(B)** Representative immunoblot of BLIMP1 expression following 48h induction of RP cells expressing *NTmiR*, *PmiR1* or *PmiR2*. **(C)** Immunofluorescence staining of BLIMP1 following 48h induction in MW cells expressing *NTmiR* or *PmiR1*, with CellProfiler quantification. **(D)** Percentage of live cells as determined by Trypan blue exclusion assay in the RP and MW cell lines comparing *PmiR1* or *PmiR2* to *NTmiR* following 48h of induction. **(E)** Immunoblot depicting lentiviral ectopic expression of EGFP or BLIMP1-EGFP in the RP *PmiR1* cells, next to the RP *NTmiR* cells. **(F)** The percentage of live RP *PmiR1* cells with ectopic EGFP or BLIMP1-EGFP expression determined by the Trypan blue exclusion assay normalised to the percentage of live cells following transduction of RP *NTmiR* cells with EGFP or BLIMP1-EGFP. BLIMP1-EGFP compared to EGFP at Day 2. **(G)** Per cent reduction as measured by resazurin assay for RP *PmiR1* cells with EGFP compared to BLIMP1-EGFP, five days after miR induction. All p-values as determined by student’s two-tailed t-test. \**p* ≤ 0.05; \*\**p* ≤ 0.01; *** *p* ≤ 0.001; \*\*\*\**p* ≤ 0.0001. All graphs were plotted as the mean of three independent experiments, with error bars representing standard deviation.

To knock-down (KD) BLIMP1, we engineered RP cells with two distinct doxycycline (dox)-inducible artificial miRNAs targeting *PRDM1* mRNA (*PmiR1 and PmiR2)*, or a non-targeting control miRNA (*NTmiR*), and MW cells with *NTmiR and PmiR1.* The induction of *PmiR1* and *PmiR2* led to the loss of BLIMP1 protein in RP cells (Fig. 1B) and *PmiR1* led to a 60% reduction in MW cells (Fig. 1C). The KD resulted in decreased cell survival, with 58% and 72% live cells remaining relative to *NTmiR* in RP *PmiR1* and *PmiR2* cell cultures respectively, 48 h post dox addition (Fig. 1D). No viable cells remained 6 days after induction of *PmiR1* or *PmiR2* (Fig. S1A). MW *PmiR1* cells also displayed an initial decrease in viability to 71% of the *NTmiR* (Fig. 1D), but by day 5 had recovered their viability, perhaps due to the survival of the non-BLIMP1 expressing cellular compartment. The proportion of apoptotic cells increased in RP cells by 2.6- and 2.3-fold 48h post *PmiR1* and *PmiR2* induction respectively (Fig. S1B). Although we consistently saw an increase in Annexin-V positive MW cells upon BLIMP1-KD, it was not statistically significant (Fig. S1C).

Crucially, the decreased cell viability upon BLIMP1-KD in RP cells was rescued in *PmiR1* cells transduced with miR-resistant BLIMP1 from a lentiviral construct [46] (Fig. 1E). BLIMP1 overexpression resulted in 81% viability, relative to BLIMP1-transduced *NTmiR* control, whereas EGFP-transduced cells were only 65% viable at 48 h post *PmiR1* induction (Fig. 1F). Five days post BLIMP1 KD, 57% of the BLIMP1-transduced cells were viable, compared to only 3% of the EGFP-transduced cells (Fig. 1F), further reflected in an increased reduction capacity of the BLIMP1 complemented cells (23% vs. 6% resazurin reduction) (Fig. 1G). Taken together, BLIMP1 is a survival factor in WM cells, likely due to the suppression of apoptosis.

### BLIMP1 expression maintains EZH2 protein levels

As studies of mouse plasmablasts and germ cells have shown a functional overlap and direct interaction of BLIMP1 and EZH2 [17, 33, 34], we investigated the potential interplay between the two factors in WM and MM. Our first line of inquiry revealed a decrease in EZH2 protein expression upon BLIMP1 KD in RP cells (Fig. 2A) and in OPM-2 MM cells also engineered with inducible *PmiR1* (Fig. 2B). Genetic complementation of miR expression with miR-resistant BLIMP1 (Fig. 1E) restored EZH2 protein levels in RP cells, confirming the effect is specifically due to BLIMP1 depletion (Fig. 2C, quantified in Fig. S2A). Furthermore, BLIMP1 and EZH2 levels were positively correlated in two separate complementation experiments (R^2^ = 0.338, R^2^ = 0.684) based on quantitation of nuclear fluorescent signal. Taken together these data reveal for the first time, a dependency of EZH2 expression on BLIMP1.

**Figure 2:**
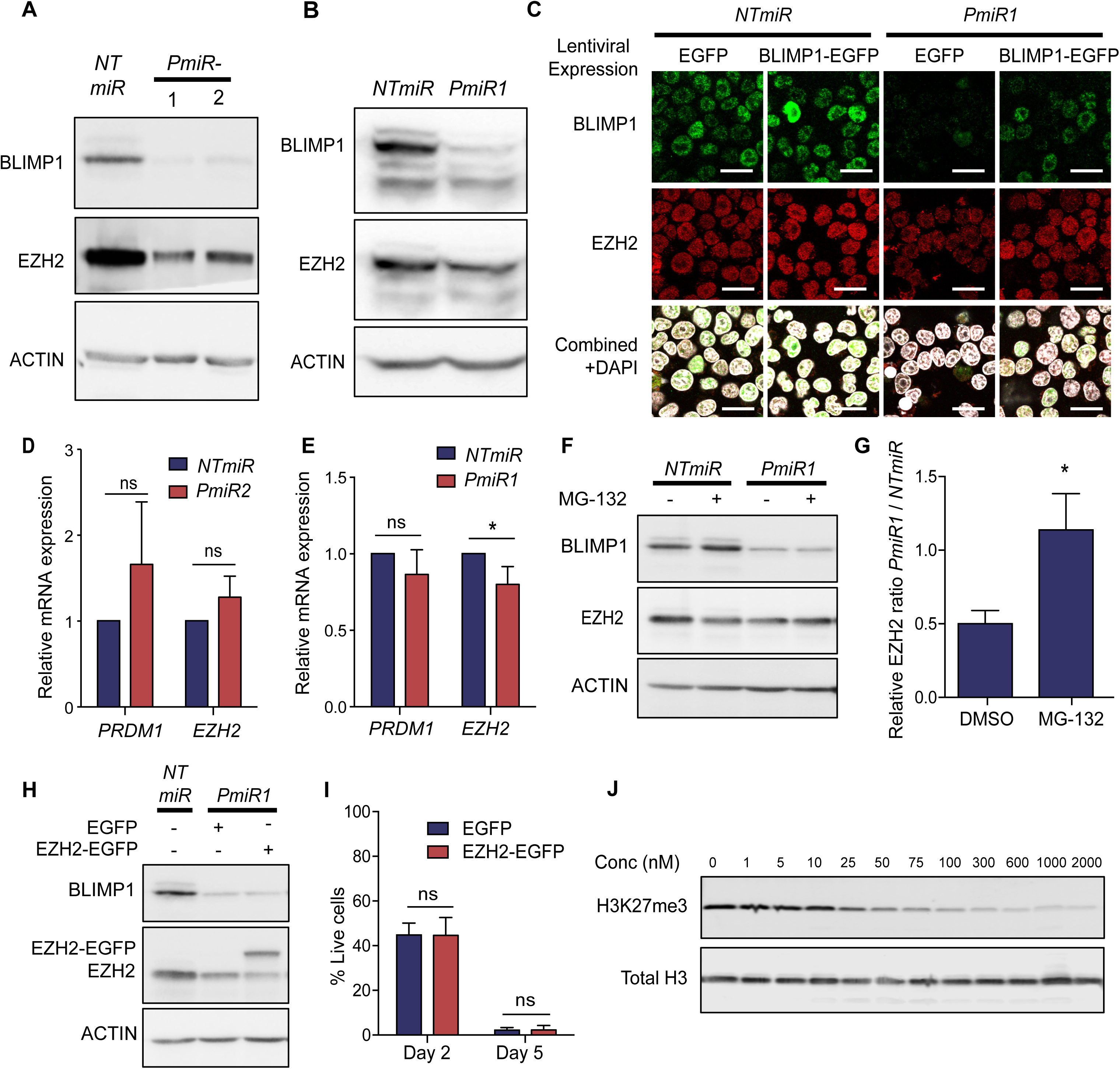
BLIMP1 maintains EZH2 protein levels in WM cells. **(A)** Immunoblot of BLIMP1 and EZH2 expression following 48h induction of RP cells expressing *NTmiR*, *PmiR1* or *PmiR2.* **(B)** Immunoblot of BLIMP1 and EZH2 expression following 48h induction of OPM-2 cells expressing *NTmiR* compared to *PmiR1*. **(C)** Immunofluorescence staining of BLIMP1 and EZH2 expression following 48 h induction of RP cells expressing *NTmiR* or *PmiR1*, transduced with EGFP or BLIMP1-EGFP. Scale bars represent 20 µm. **(D)** RT-qPCR results depicting relative mRNA expression of *PRDM1* and *EZH2* normalised to *PPIA* and *ACTB* in the RP cell line and **(E)** in the OPM-2 cell line. **(F)** Immunoblot of BLIMP1 and EZH2 expression following 24 h induction of RP cells expressing *NTmiR* or *PmiR1* treated with DMSO or 5µM MG-132 for 4h. **(G)** The ratio of EZH2 expression relative to actin for RP *PmiR1* cells divided by RP *NTmiR* cells treated with DMSO or MG132 as in Fig. 2F. **(H)** Immunoblot of BLIMP1 and EZH2 expression following 48 h induction of RP *NTmiR* cells or *PmiR1* cells transduced with EGFP or EZH2-EGFP. **(I)** Percentage of live cells as determined by Trypan blue exclusion assay in the RP *PmiR1* cells transduced with EGFP or EZH2-EGFP. All p-values as determined by student’s two-tailed t-test. (ns) *p* > 0.05; \**p* ≤ 0.05; \*\**p* ≤ 0.01; *** *p* ≤ 0.001; \*\*\*\**p* ≤ 0.0001. All graphs were plotted as the mean of three independent experiments with error bars representing standard deviation. **(J)** Immunoblot of histone extracts from RP cells stained for H3K27me3 with total H3 as a loading control following 48h tazemetostat treatment at the indicated concentrations. The 0 concentration was treated with vehicle control, DMSO.

Reverse transcription followed by quantitative PCR (RT-qPCR) revealed that *EZH2* mRNA levels were unchanged in RP *PmiR2* cells upon BLIMP1 KD and only slightly decreased in OPM-2 cells upon *PmiR1* induction, indicating that the loss of EZH2 expression is at the post-transcriptional level (Fig. 2D-E). Notably, the treatment of RP *PmiR1* cells with the proteasome inhibitor MG-132, restored EZH2 to the same level as that of *NTmiR* cells also treated with proteasome inhibitor (Fig. 2F-G), revealing that BLIMP1 modulates EZH2 by inhibition of proteasome mediated degradation.

However, complementing the RP *PmiR1* cells with ectopic EZH2 expression (Fig. 2H) failed to increase their viability (Fig. 2I). Consistent with this, the EZH2-specific catalytic inhibitor tazemetostat did not affect RP cell viability even over a 96 h period (Fig. S2B), despite a dose dependent decrease in H3K27me3 levels (Fig. 2J), nor did 7 days of treatment. Taken together, BLIMP1 maintains EZH2 protein levels via modulation of proteasome mediated degradation, but the effect of BLIMP1 on cell survival is independent of EZH2.

### BLIMP1 KD induces large transcriptional changes

To investigate the overlap of BLIMP1 and EZH2 in downstream gene regulation in WM cells, we performed transcriptome profiling of the RP *PmiR2* and *NTmiR* cells following 48 h of induction. Using a q-value cutoff of 0.05, we identified 7814 differentially expressed genes between *PmiR2* and *NTmiR* (Fig. 3A and Table S1).

**Figure 3:**
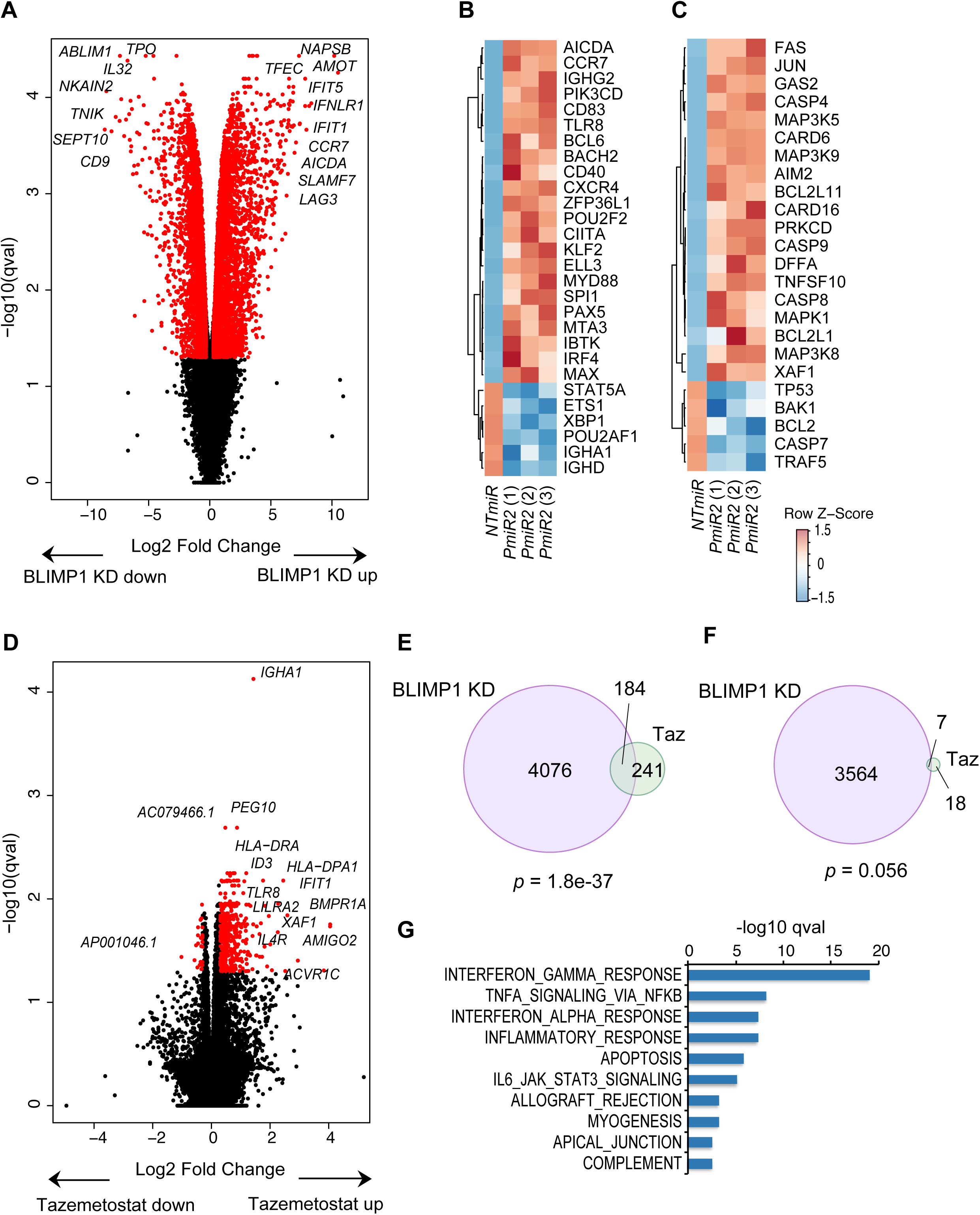
BLIMP1 KD and EZH2 inhibition induce overlapping transcriptional changes. RNAseq results for **(A)** 48h-induced RP cells comparing *PmiR2* to *NTmiR*. Values are plotted as log_2_ fold change vs. –log_10_(q-value). Red indicates those genes with a q-value ≤ 0.05 and a log_2_ fold change ≤ −0.3 or ≥ 0.3. Heat maps depicting the Z-score of the log_2_ fold change comparing *PmiR2* to *NTmiR* for three independent replicates looking at **(B)** B cell genes and **(C)** apoptosis genes. **(D)** RNAseq results for RP cells treated for 48h with 300nM tazemetostat compared to vehicle control (DMSO). **(E)** Overlapping genes with increased expression following *PRDM1* KD or tazemetostat treatment. **(F)** Overlapping genes with decreased expression following *PRDM1* KD or tazemetostat treatment. Overlaps tested using Fisher’s exact test. **(G)** Overlaps of genes significantly induced by both BLIMP1 KD and tazemetostat with Hallmarks gene sets from the molecular signatures database, showing the top 10 most significantly overlapping gene sets.

Consistent with BLIMP1 repressing the B cell transcriptional program [15, 17, 19, 47], previously characterised B cell lineage targets of BLIMP1, including *CIITA* [48, 49], *PAX5* [50], *SPIB* and *BCL6* [15] were repressed by BLIMP1 in our study (Fig. 3B). However, other BLIMP1 targets such as *MYC* [51] and *ID3* [15] were unaltered. Curiously, the myeloma-driving transcription factor *IRF4* [44], which is activated downstream of BLIMP1 in plasma cells [17] was repressed by BLIMP1 in RP cells (Fig. 3B). Interestingly, the BTK inhibitor, *IBTK* was also repressed by BLIMP1. The inhibition of BTK via Ibrutinib is now a common treatment for WM [52]. The expression of key apoptosis genes was increased following BLIMP1 KD (Fig. 3C) and finally, we also observed a de-repression of *SMURF2* mRNA (Fig. S3A). SMURF2 is an E3 ubiquitin ligase known to target EZH2 for proteasome-mediated degradation and could therefore contribute to the decrease in EZH2 protein levels upon the loss of BLIMP1 [53]. Taken together, BLIMP1 KD induces extensive gene expression changes in RP cells including the de-repression of B cell- and apoptosis-related genes, as well as *SMURF2*.

### BLIMP1 and EZH2 regulate overlapping pathways

Because of the dependency of EZH2 on BLIMP1 levels, we wanted to determine to which extent the effect of BLIMP1 on transcription is dependent on EZH2’s catalytic activity, and therefore performed transcriptome profiling upon catalytic inhibition of EZH2 with tazemetostat. This resulted in 450 differentially expressed genes compared to vehicle alone (Fig. 3D and Table S2). The amplitude of change for individual genes was smaller than the changes induced by BLIMP1 KD (Comparing Fig. 3D and 3A). Nevertheless, a highly significant overlap emerged between transcripts increased in the BLIMP1 KD and tazemetostat treated cells (184 genes, p = 1.8e-37) (Fig. 3E, Table S3), but the genes with decreased expression did not significantly overlap (Fig. 3F, Table S4). Tazemetostat led to the de-repression of B cell identity genes, including *STAT5B* and *ID3* (Fig. S3B) and interestingly, despite EZH2 inhibition not affecting cell survival, a number of apoptosis genes were differentially expressed, including *XAF1*, *CASP4*, *FAS* and *JUN* (Fig. S3C), suggesting a possible sensitisation to apoptosis.

To investigate the pathways jointly regulated by BLIMP1 and EZH2, we performed an overlap analysis using the molecular signatures database [54, 55] of the 184 genes induced by both BLIMP1 KD and tazemetostat. Several gene sets were significantly enriched, including interferon and TNFα responses, the inflammatory response and apoptosis (Fig. 3G), consistent with the known roles of BLIMP1 [14, 18, 56–58], and suggesting that pathways involved in immune responses are regulated in concert by BLIMP1 and EZH2. Collectively, the transcriptomic analyses demonstrate a large overlap in targets of repression by BLIMP1 and EZH2, highlighting the interplay of the two factors.

### BLIMP1 binds to a set of H3K27me3 marked genes at a distance from the mark

Given the overlap in genome wide positioning of BLIMP1 and the H3K27me3 mark in mouse plasmablasts [17], we examined whether the overlapping effects of BLIMP1 and EZH2 on gene transcription could be modulated not only through the regulation of EZH2 by BLIMP1 but also by BLIMP1 potentially recruiting EZH2 to chromatin in WM cells. We therefore performed chromatin immunoprecipitation coupled to deep sequencing (ChIPseq) for H3K27me3 and BLIMP1 in RP cells. We identified 14946 H3K27me3 peaks (Table S5), assigned to 4198 genes (Table S6), and 506 BLIMP1 peaks (Table S7), assigned to 841 genes (Table S8). If BLIMP1 recruits EZH2 to chromatin in WM cells, a large proportion of BLIMP1 peaks would be located in close proximity to H3K27me3 marks. However, only 8 sites bore both H3K27me3 and BLIMP1 within 500 bp (Fig. 4A) and only a small level of bimodel enrichment of H2K27me3 was observed in the ± 3 kb flanking BLIMP1 peaks (Fig. 4B). Interestingly however, when we compared the genes assigned to the peaks in the respective experiments, 261 genes were both bound BLIMP1 and marked by H3K27me3 in RP cells, a highly statistically significant overlap (*p* = 2.9e-25) (Fig. 4C, Table S9). An analysis [54, 55] of the genes marked by both BLIMP1 and H3K27me3 in the RP cells demonstrated significant enrichment for the same immune signalling gene sets as in the RNAseq data, as well as genes relating to the estrogen response, hypoxia and the p53 pathway (Fig. S4A).

**Figure 4:**
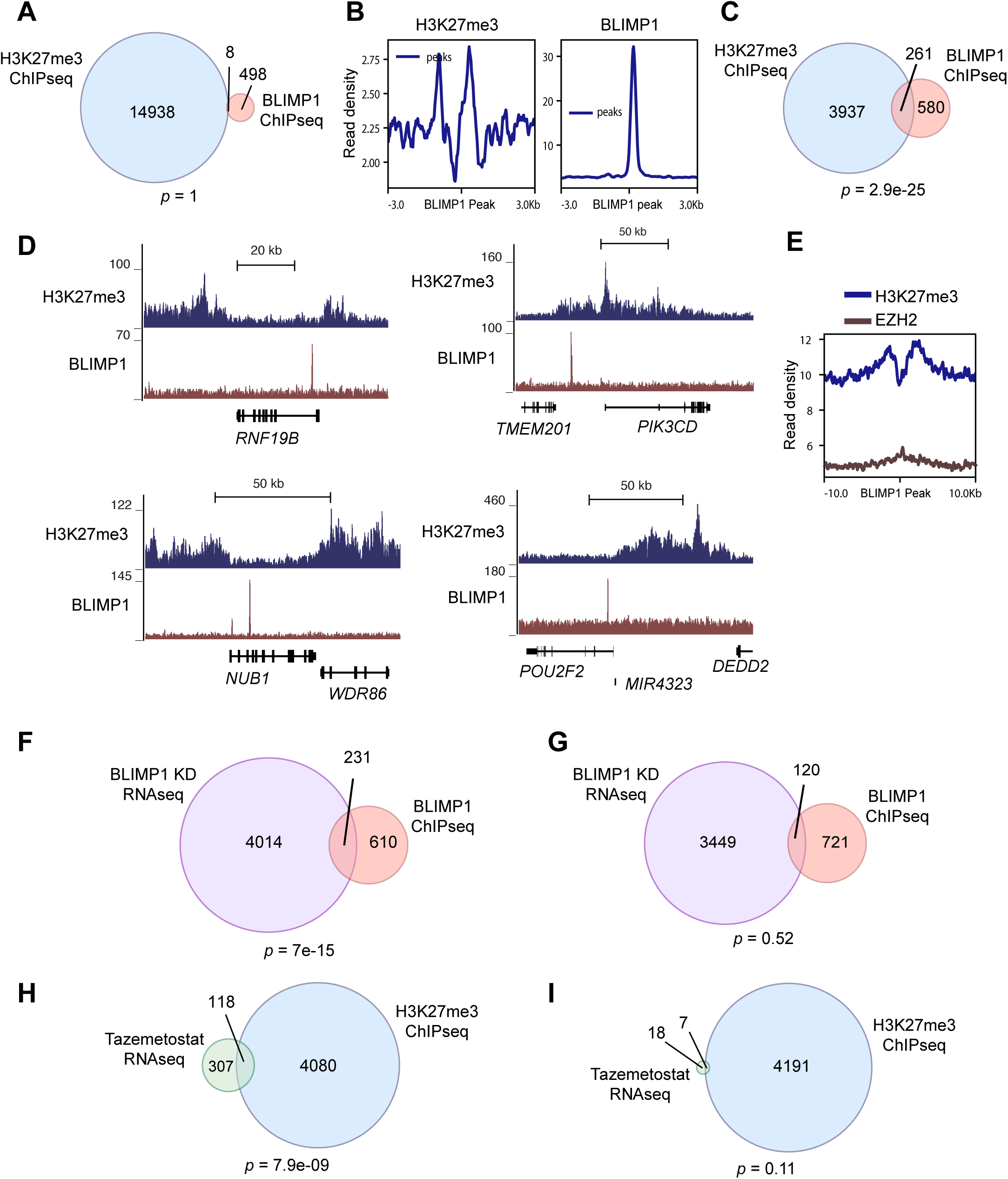
BLIMP1 binds at a distance to the H3K27me3 mark. **(A)** Venn diagram of H3K27me3 and BLIMP1 peaks extended ±500 bp, showing overlaps in these regions. (ns) not significant as determined by hypergeometric test. Called peaks determined by overlap from peak calling from two independent experiments. **(B)** Enrichment of signal from ChIPseq tracks ±3kb from the centre of BLIMP1 binding sites. Data depicts representative experiment of two biological replicates. **(C)** Venn diagram of genes assigned to H3K27me3 and BLIMP1 peaks showing overlapping genes. p= 2.9e-25 as determined by Fisher’s exact test. **(D)** ChIPseq tracks for H3K27me3 and BLIMP1 in the RP cell line. Data represents combination of reads from two independent experiments. **(E)** Enrichment of signal from ChIPseq tracks ±10kb from the centre of BLIMP1 binding sites in the NCI-H929 cell line. **(F)** Venn diagram depicting the overlap in genes with significantly increased expression following BLIMP1 KD and genes assigned to BLIMP1 binding sites. p= 7e-15, as determined by Fisher’s exact test. **(G)** Venn diagram depicting the overlap in genes with significantly decreased expression following BLIMP1 KD and genes assigned to BLIMP1 binding sites. (ns) not significant as determined by Fisher’s exact test. **(H)** Venn diagram depicting genes with significantly increased expression following tazemetostat treatment overlapping with genes assigned to H3K27me3 peaks. p= 7.9e-9, as determined by Fisher’s exact test. **(I)** Venn diagram depicting genes with significantly decreased expression following tazemetostat treatment overlapping with genes assigned to H3K27me3 peaks. (ns) not significant, as determined by Fisher’s exact test. All ChIPseq experiments were performed as two biological replicates.

To test whether these findings apply also to MM, we performed ChIPseq for BLIMP1 and H3K27me3 in the OPM-2 (Tables S10-13) and NCI-H929 (Tables S14-17) myeloma cell lines, and EZH2 in NCI-H929 cells (Tables S18-19). As with the RP cells, we rarely observed peaks within a 500bp distance when comparing BLIMP1 and H3K27me3 (Fig. S4B) or BLIMP1 and EZH2 (Fig. S4C) in the OPM-2 and NCI-H929 cell lines. However, we observed a statistically significant overlap between genes bound by BLIMP1 and H3K27me3 in NCI-H929 cells, but not OPM-2 cells (Fig. S4D). Genes bound both by BLIMP1 and EZH2 were not statistically over-represented in NCI-H929 cells (Fig. S4E). The above analyses reveal that the majority of BLIMP1 and EZH2 binding to chromatin occurs at relatively distant sites both in MM and WM cells. Thus, their direct regulation on chromatin is unlikely to be due to a direct physical interaction.

The known DNA binding motif for BLIMP1 [59] was enriched by *de novo* motif analysis in the BLIMP1 peaks for all three cell lines (Fig. S4F) and the BLIMP1 ChIPseq signals were enriched at proximal positions relative to their assigned transcription start sites (TSSs) (Fig. S4G). Comparing the distribution of BLIMP1 peaks in OPM-2 and NCI-H929 to the RP cell line (Fig. S4H), showed that BLIMP1 binds to largely the same sites in WM and MM.

An analysis [54, 55] of genes assigned to H3K27me3 peaks for each cell line revealed similar gene sets marked by H3K27me3 in all three cell lines (Fig.ure S5A). Interestingly, TNFα and IL2/STAT5 signalling genes were amongst the most highly enriched in the RP but not the myeloma cells. The myeloma cell lines displayed a much stronger enrichment of H3K27me3 over TSSs than the RP cells (Fig. S5B), indicating that H3K27me3 is more often present at gene distal sites in RP cells than myeloma cells. EZH2 was also highly enriched just downstream of TSSs in NCI-H929 cells (Fig. S5C). Correspondingly, we observed low enrichment of the H3K27me3 mark in the OPM-2 and NCI-H929 cell lines over sites marked by H3K27me3 in the RP cell line (Fig. S5D). Interestingly, *SMURF2* which can affect EZH2 stability [53], was bound by BLIMP1 in all three cell lines and bore the H3K27me3 mark in the RP cells as well as EZH2 in the NCI-H929 cell line (Fig. S5E).

To identify the direct transcriptional targets of BLIMP1 and EZH2, we compared our ChIPseq and transcriptome profiling data from the RP cell line. There were 231 and 120 genes associated with BLIMP1 binding that were either induced or repressed upon BLIMP1 KD respectively (Fig. 4F-G). Conversely, 118 genes were both induced upon tazemetostat treatment and associated with the H3K27me3 mark (Fig. 4H), but only 7 genes with decreased expression were marked by H3K27me3 (Fig. 4I). Overall, only 3% of genes marked by H3K27me3 were de-repressed following tazemetostat treatment in these experiments, indicating that inhibition of EZH2’s catalytic activity alone is insufficient to activate most H3K27me3 targets over 48 h. Meanwhile, approximately one third of BLIMP1-bound genes showed altered expression upon BLIMP1 KD, indicating that BLIMP1 binding actively maintains gene repression. Taken together, as BLIMP1 and H3K27me3 are largely present at distinct sites from one another, and most of their overlapping effects on transcription are likely through binding to the same genes at distinct sites, as well as BLIMP1 maintenance of EZH2 protein levels.

### A subset of BLIMP1 targets are regulated via EZH2

A number of genes repressed by BLIMP1 bore both BLIMP1 and the H3K27me3 mark, such as *PIK3CD*, *POU2F2* (Fig. 4D) and *ZFP36L1* (Fig. 5A). Meanwhile, others bore only the H3K27me3 mark, but not BLIMP1, including *RCAN3*, *TNFRSF14* and *CIITA* (Fig. 5B-D). Yet others such as the macrophage gene *TFEC* [60] were bound by BLIMP1 alone (Fig. 5E). We therefore asked whether the genes bearing the H3K27me3 mark are repressed by BLIMP1 via the maintenance of EZH2. By overexpressing EZH2 in RP cells upon BLIMP1 KD, we performed RT-qPCR and observed that repression of the genes *RCAN3* and *ZFP36L1* was restored (Fig. 5F) as well as the repression of the immune inhibitory checkpoint ligand/receptor gene *TNFRSF14* [61] and the inhibitory receptor gene *HAVCR2* [62]. By comparison, BLIMP1 binding targets not bearing the H3K27me3 mark such as *TFEC* were not altered upon EZH2 restoration. Thus, a subset of BLIMP1 targets are regulated through EZH2 maintenance and can be identified by the H3K27me3 mark, whereas others are regulated independently of EZH2.

**Figure 5:**
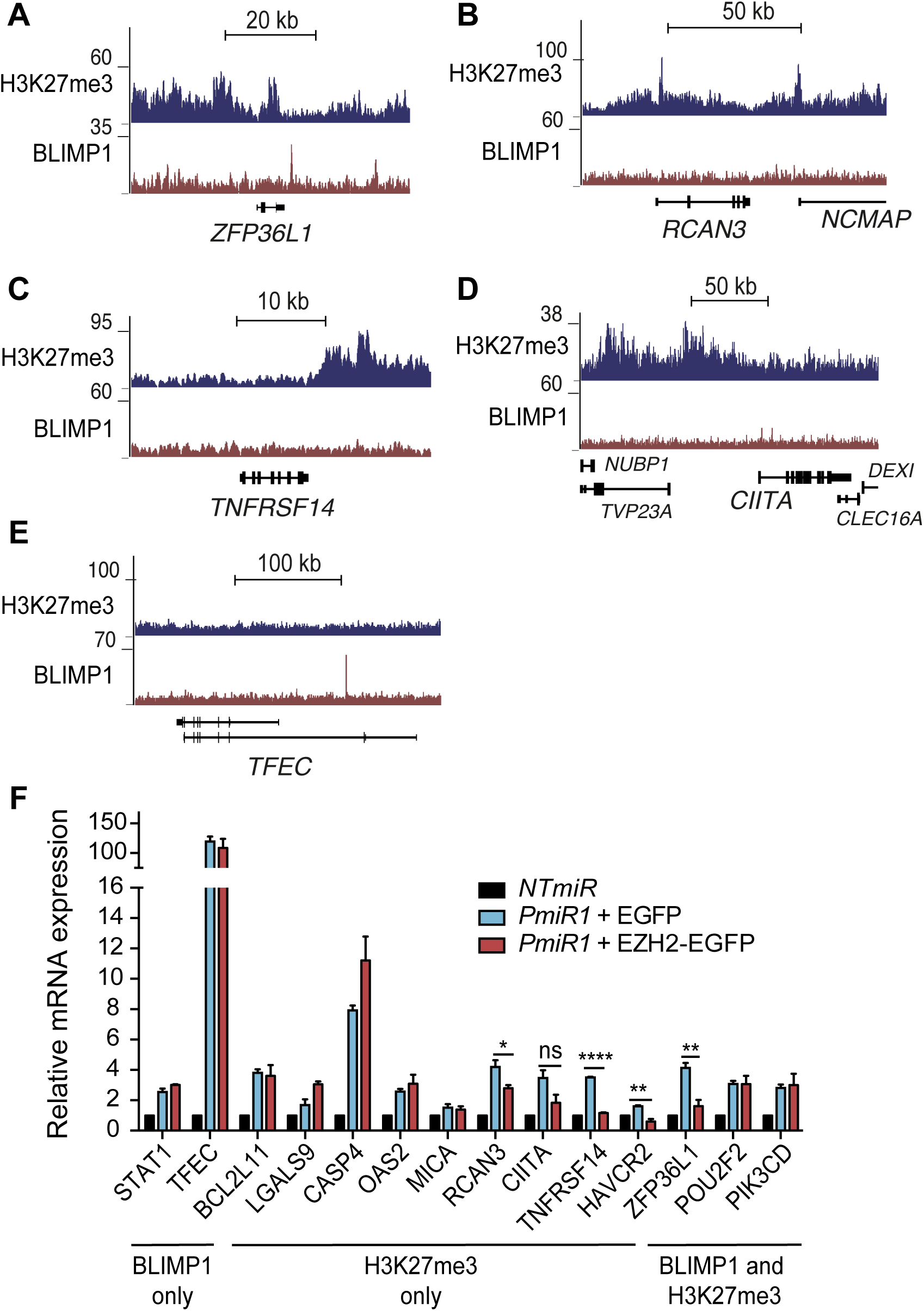
BLIMP1 regulates a subset of target genes via EZH2. ChIPseq tracks for H3K27me3 and BLIMP1 over the genes **(A)** *ZFP36L1*, **(B)** *RCAN3*, **(C)** *TNFRSF14*, **(D)** *CIITA*, and **(E)** *TFEC*. **(F)** Bar graph depicting RT-qPCR experiments in RP cells expressing *NTmiR* or *PmiR1* with EGFP or EZH2-EGFP. The selected target genes bear peaks for either BLIMP1, H3K27me3 or both factors. All p-values as determined by student’s two-tailed t-test. (ns) *p* > 0.05; \**p* ≤ 0.05; \*\**p* ≤ 0.01; \*\*\*\**p* ≤ 0.0001. The bar graph was plotted as the mean of three independent experiments with error bars representing standard deviation.

### BLIMP1 represses transcription of immune surveillance and signalling molecules in concert with EZH2

Further analysis of the highly enriched immune signalling genes differentially expressed upon BLIMP1 KD identified three mechanistic categories. First, BLIMP1 represses the expression of genes encoding surface ligands that can activate T and NK cells, including *ICOSLG*, *TNFSF9*, *CD48*, *MICA*, *CLEC2B*, *ICAM1* and *ITGAM*, and MHC class II molecules including *HLA-DMA*, *HLA-DMB* and *CD1D* (Fig. 6A). Second, BLIMP1 represses genes encoding both immune-checkpoint inhibitory ligands and their corresponding receptors, including the recptor-ligand pairs *TNFRSF14* and *BTLA*, as well as *LAG3* and *LGALS3*. Furthermore, the inhibitory receptor gene *LILRB1,* whose loss promotes immune escape in myeloma [63] is repressed by BLIMP1. Additionally, the gene encoding the inhibitory ligand PD-L2 is repressed, with BLIMP1 binding just downstream of *PDCDLG2* (Fig. S6A). Notably, repression of inhibitory ligands and receptors could promote or inhibit escape from immune surveillance. The third mechanistic category includes interferon and TNFα signalling genes. The receptor-encoding genes *IFNGR1*, *IFNAR1*, *IFNLR1*, *TNFRSF1A* and *TNFRSF1B* are de-repressed upon BLIMP1 KD, as well as downstream players in interferon signalling, *JAK1*, *STAT1*, *STAT2*, *IRF9 IFIT1-3*, *OAS1-3*, and TNF pathway members *MAP3K5*, *CASP8* and *CASP9*, which are also apoptosis mediators (Fig. 3C). Beyond these categories, MHC class I genes, *HLA-A* and *HLA-B* which can both mediate inhibition of NK-cell immune surveillance, are induced by BLIMP1, in contrast to previous studies [58, 64].

**Figure 6:**
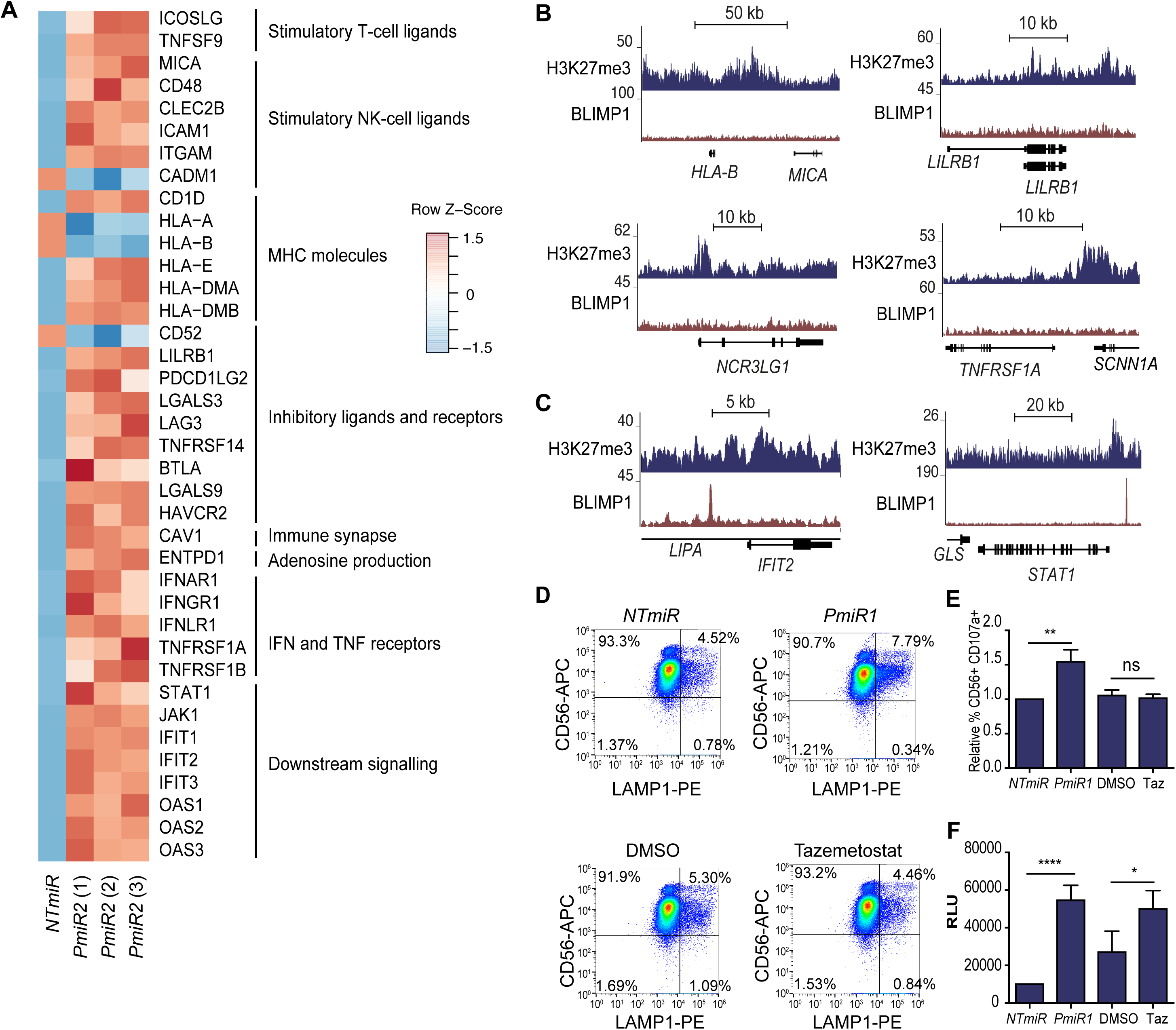
BLIMP1 and EZH2 promote WM tumour immune evasion. **(A)** Heat maps showing z-score of the log2 fold change for *PmiR2* compared to *NTmiR* and in RP cells with three independent replicates looking at genes involved in stimulation of T and NK cells, MHC molecules, inhibitory ligands and receptors, IFN and TNF receptors and downstream signalling. ChIPseq tracks for H3K27me3 and BLIMP1 in the RP cell line over **(B)** the immune surface molecule genes *HLA-B, MICA, LILRB1, NCR3LG1, TNFRSF1A*, or **(C)** the downstream signalling genes, *IFIT2* and *STAT1*. Data represents a combination of reads from two independent experiments. **(D)** Percentage CD56^+^LAMP1^+^ cells (Q2) representing degranulating NK cells, as determined by flow cytometry following co-culture with RP cells with *NTmiR* or *PmiR1*, or RP cells treated with DMSO or Tazemetostat. One representative experiment is displayed. **(E)** Relative quantification of the degranulation assay. **(F)** Cytotoxicity depicted in relative luminescence units (RLU), as measured by adenylate kinase activity in the culture media following 4 h co-culture of NK cells with RP cells expressing *NTmiR* or *PmiR1*, or RP cells treated with DMSO or 1µM Tazemetostat. Cells were co-cultured at the effector:target ratio of 20:1. Results of the degranulation and cytotoxicity assays from four individual donors, performed in two pairs on two separate occasions. All p-values as determined by student’s two-tailed t-test. (ns) *p* > 0.05; \**p* ≤ 0.05; \*\**p* ≤ 0.01; \*\*\*\**p* ≤ 0.0001. All graphs were plotted as the mean of three independent experiments with error bars representing standard deviation.

Around half of the genes discussed above were also significantly changed with tazemetostat treatment (Fig. S6B), including *CD48*, *LILRB1*, *LGALS3, PDCD1LG2* and *STAT1*. Interestingly, while none of the surface molecule genes except for *PDCDLG2* were bound by BLIMP1, some were enriched for H3K27me3 (Fig. 6B). Whereas the downstream signalling effectors, *IFIT2* and *STAT1*, were bound by BLIMP1 (Fig. 6C), consistent with previous findings [65]. In addition, a number of these differentially expressed genes did not bear either H3K27me3 or BLIMP1, and their expression changes are therefore likely secondary effects downstream of EZH2 and BLIMP1. Taken together, BLIMP1 and EZH2 repress the transcription of genes that encode mediators of killing by NK or T cells when expressed on target cells, NK- and T-cell inhibitory receptors and their ligands, as well as cytokine receptors and downstream signal propagation.

### BLIMP1 and EZH2 confer evasion from NK cell-mediated cytotoxicity

The transcriptional changes above suggested that BLIMP1 and EZH2 were potentially mediating escape from tumour immune surveillance [66]. We therefore hypothesised that the alterations in gene expression upon the loss of BLIMP1 protein or EZH2 activity could lead to changes in NK cell-mediated cytotoxicity. We isolated NK cells from human blood and co-cultured them with RP cells after BLIMP1 KD or treatment with tazemetostat and measured the surface expression of the degranulation marker LAMP-1 (also called CD107a) on NK cells by flow cytometry. This revealed a 1.5-fold increase in frequency of LAMP-1^+^ NK cells upon co-culture with *PmiR1* compared to *NTmiR* RP cells, showing that the loss of BLIMP1 sensitises NK-cells to activation by RP (Fig. 6D-E and S5D), however tazemetostat-treated RP cells did not increase NK-cell degranulation. Strikingly though, both BLIMP1 KD and tazemetostat treatment resulted in increased NK cell-mediated death of RP cells (Fig. 6F). Collectively, these data indicate that BLIMP1 and EZH2 drive the escape from NK-cell immune surveillance in WM, with BLIMP1 both suppressing NK cell activation and the resultant WM cell death, whereas EZH2 suppresses the cell death response of WM cells.

## Discussion

In this study we show for the first time that the plasma cell master regulator BLIMP1 has a crucial role in WM cell survival and maintains the protein levels of the histone methyltransferase EZH2, demonstrating a more complex functional interaction between BLIMP1 and EZH2 than previously thought [17, 32–34]. Furthermore, we show that the factors mediate the evasion of immune surveillance, as evidenced by the enhanced degranulation of NK cells upon loss of BLIMP1 in WM cells, and enhanced NK cell-mediated WM cellular cytotoxicity upon either BLIMP1 KD or EZH2 inhibition.

Our finding that BLIMP1 promotes survival of WM cells provides new insight into WM biology and potentially its aetiology. BLIMP1 is activated downstream of MYD88 signalling in B cells [29], and BLIMP1 expression is increased in tumours harbouring the MYD88^L265P^ activating mutation in WM [30]. However, heterozygous deletions in the 6q locus containing *PRDM1* are increased in MYD88^L265P^ tumours [24]. As high levels of BLIMP1 inhibit proliferation, and intermediate levels promote survival and immunoglobulin (Ig) secretion [47, 67], our findings might imply that the loss of one copy of the *PRDM1* gene subsequent to MYD88 constitutive activation dampens the anti-proliferative effect of BLIMP1 while still maintaining its positive effect on survival and Ig secretion. Indeed, we observed decreased cell viability upon the loss of BLIMP1 even in MW cells that expresses low BLIMP1 levels. A complete loss of BLIMP1 during tumorigenesis has been shown to give rise to B cell lymphomas in mice [21–23], but the loss of Ig secretion ability and plasma cell differentiation would likely preclude WM tumour formation, with the subsequent requirement for BLIMP1 in survival.

A first line of treatment in WM is rituximab therapy, targeting the B cell specific surface molecule CD20 at least in part through NK-cell engagement [68]. However, the plasma cells remaining after this treatment present one of the biggest challenges in WM therapy [69]. Futhermore, rituximab is not recommended for patients exhibiting high serum IgM levels [70], due to the risks of an IgM ‘flare’ following treatment [71]. This highlights the importance of the antibody-secreting CD20^low^ or negative plasma cell compartment in WM [69, 72], which is likely maintained by BLIMP1.

Interestingly, the requirement for BLIMP1 in mediating WM cell survival is through other mechanisms than its effect on EZH2. While both MM and normal plasma cells rely on BLIMP1 for their survival [18, 19, 73], EZH2 expression is decreased in mature plasma cells [74, 75]. Nevertheless, many MM cell lines are reliant on EZH2 for cytokine independent growth and thus their more aggressive plasma cell leukemia phenotype [40, 75]. Our results indicate that the effects of BLIMP1 on EZH2 levels might extend to myeloma, and perhaps other tumours where the two factors are co-expressed, and warrants further study.

Immune evasion is most extensively studied in relation to solid tumours, but in lymphomas, the paradigm is somewhat different. Lymphocytes interact with other cells of the immune system not only when they are abnormal, such as in cancer, but also to receive stimulatory and inhibitory signals, regulating their own functions [76]. In WM, secreted PD-1 ligands inhibit T cell responses [77], but other mechanisms of immune evasion have not been investigated. Our results demonstrate for the first time that BLIMP1 and EZH2 mediate evasion from NK cell surveillance, both in terms of the suppression of NK cell activation downstream of BLIMP1 as well as resistance to NK cell mediated cytotoxicity downstream of both factors. Based on our transcriptomic profiling, BLIMP1 likely promotes immune evasion in WM by repressing activating ligands and MHC class I molecules leading to “hiding” from cytotoxic lymphocytes. Conversely, EZH2 is more likely to be de-sensitising WM cells to external cytotoxicity-inducing signals perhaps by repressing pro-apoptotic gene expression. Interestingly, in melanoma EZH2 promotes evasion from IFNγ-producing cytotoxic T cells [78]. This finding together with our current study make EZH2 an interesting target for chemical sensitisation of WM and other tumours to immune-cell mediated killing.

In conclusion, we provide evidence for a crucial role of BLIMP1 in promoting WM cell survival and we show that BLIMP1 maintains EZH2 protein levels in both WM and MM cells. We further reveal a cooperation between BLIMP1 and EZH2 in the repression of immune surveillance genes, and find that BLIMP1 and EZH2 confer evasion from NK-cell mediated cytotoxicity. In future studies it would be important to further investigate the intricate interplay between BLIMP1 and EZH2 in cancers to expand their therapeutic potential.

## Data availability

RNAseq and ChIPseq data are available at http://www.ebi.ac.uk/arrayexpress/experiments/E-MTAB-7739 under the accession code: E-MTAB-7739.

## Acknowledgements

We thank Arnar Palsson and Dagny A. Runarsdottir for their advice on RNAseq data analysis, Jona Freysdottir and Sunnefa Yeatman Omarsdottir for their advice on the isolation of NK cells, and the group of Eirikur Steingrimsson for useful discussions and advice on the project. We thank Dr. Roopsha Sengupta for providing critical inputs and proof-reading the manuscript.

