## Supplementary Figures for "The BLIMP1 – EZH2 nexus in a non-Hodgkin lymphoma"

#### Supplementary Figure 1

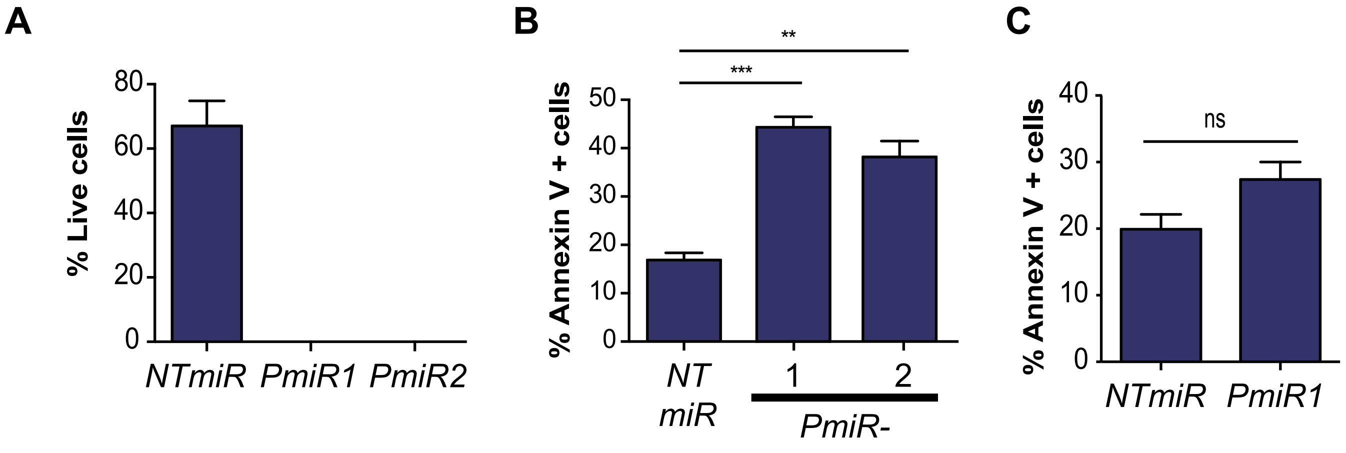


Supplementary Figure 1

(A) Percentage of live cells as determined by Trypan blue exclusion assay in the RP cells comparing PmiR1 or PmiR2 to NTmiR following 6 days of induction. (B) Annexin V staining of RP cells induced for 48h with dox to express NTmiR, PmiR1 or PmiR2; **$p \leq 0.01$; *** $p \leq 0.001$ (C) Annexin V staining of 48h-induced MW cells with NTmiR or PmiR1; $p = 0.0572$.

**Supplementary Figure 2**

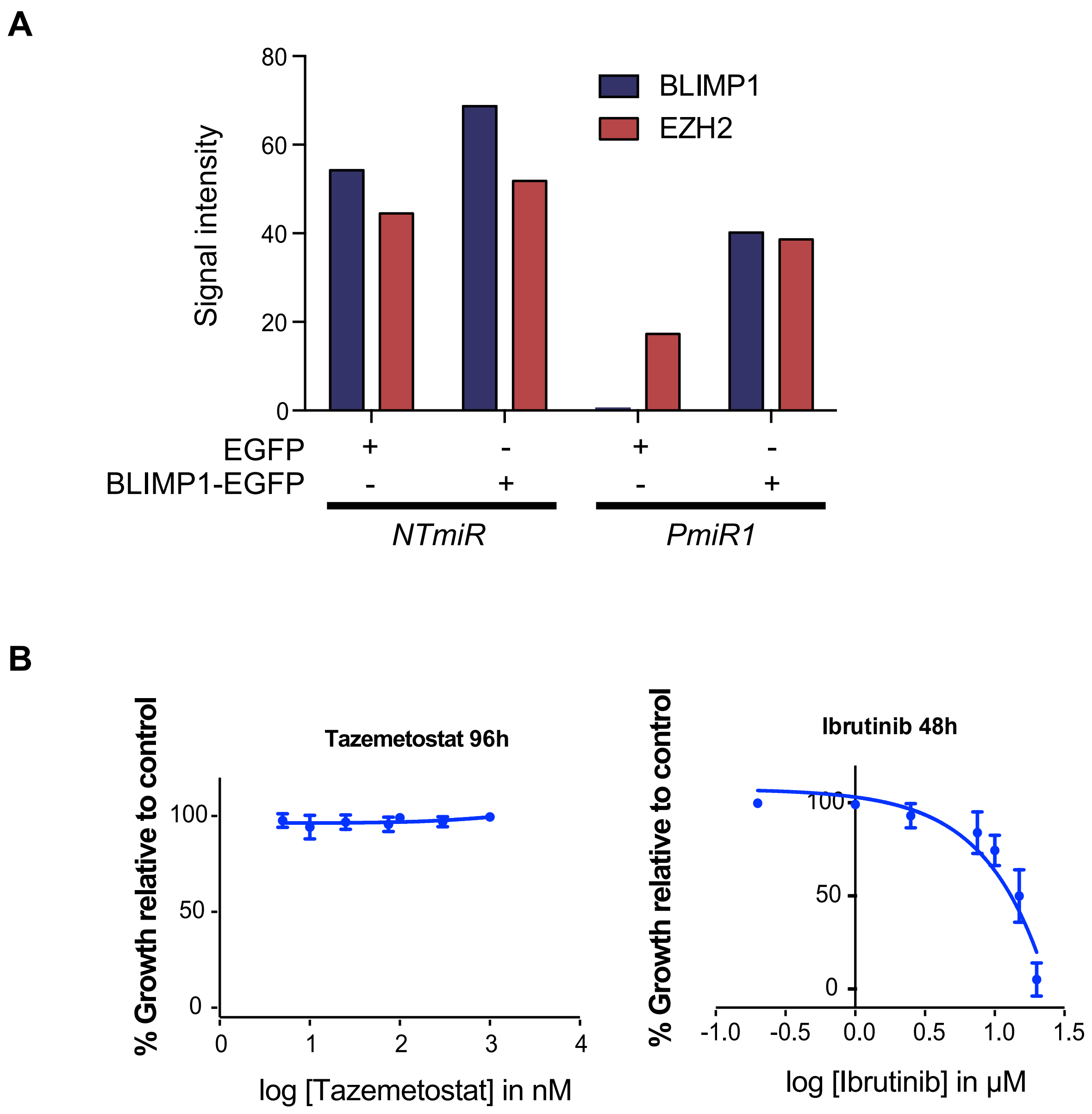


Supplementary Figure 2

(A) Signal intensity of immunofluorescence staining for BLIMP1 and EZH2 as quantified by cell profiler from approximately 5000 cells per sample across two independenct replicates. (B) Relative growth of RP cells treated with tazemetostat for 96h, shown next to RP cells treated with Ibrutinib for 48h.

#### Supplementary Figure 3

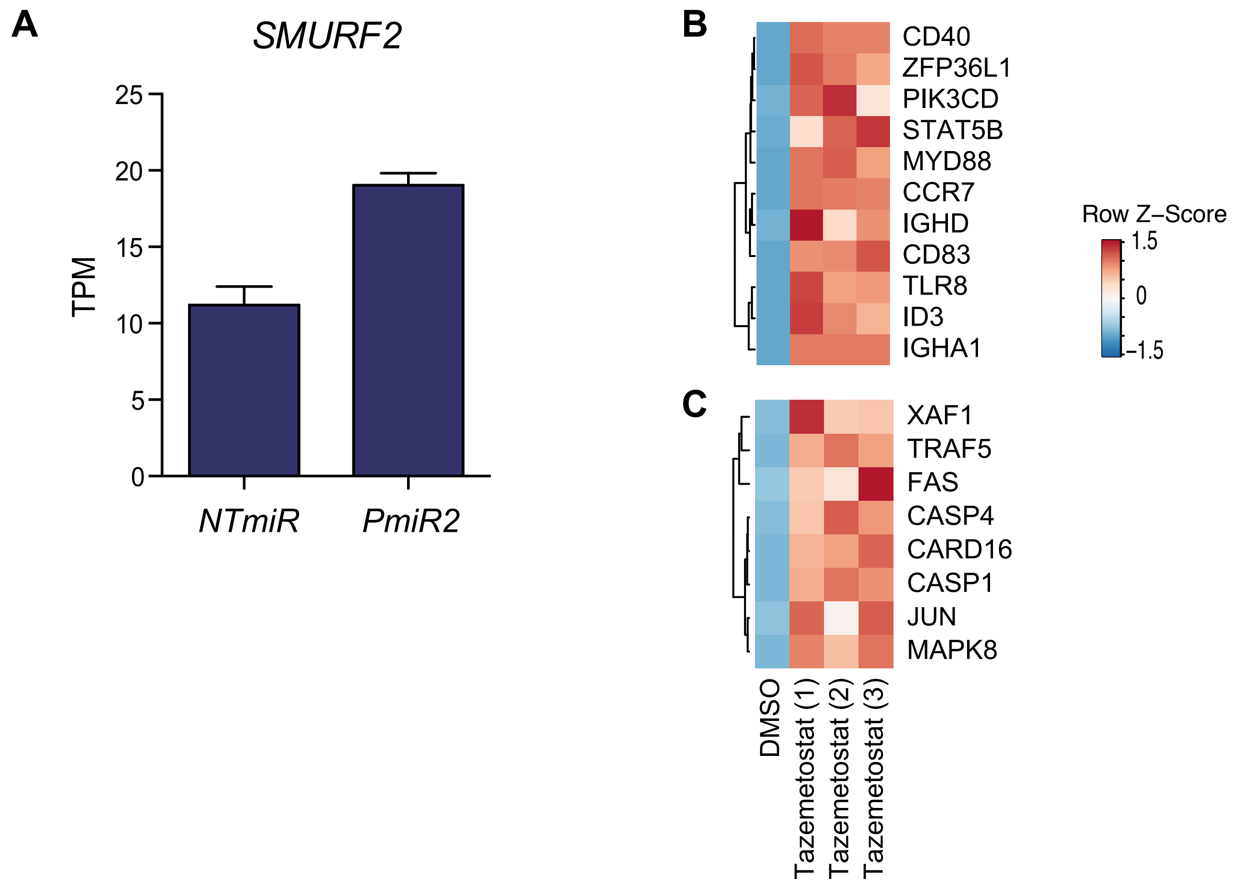


Supplementary Figure 3

(A) Graph of RNAseq results depicting normalised transcripts per million (TPM) values for SMURF2 in RP cells with NTmiR and PmiR1. Heat maps depicting the Z-score of the log2 fold change comparing tazemetostat treatment to DMSO for three independent replicates looking at (B) B cell genes and (C) apoptosis genes.

#### Supplementary Figure 4

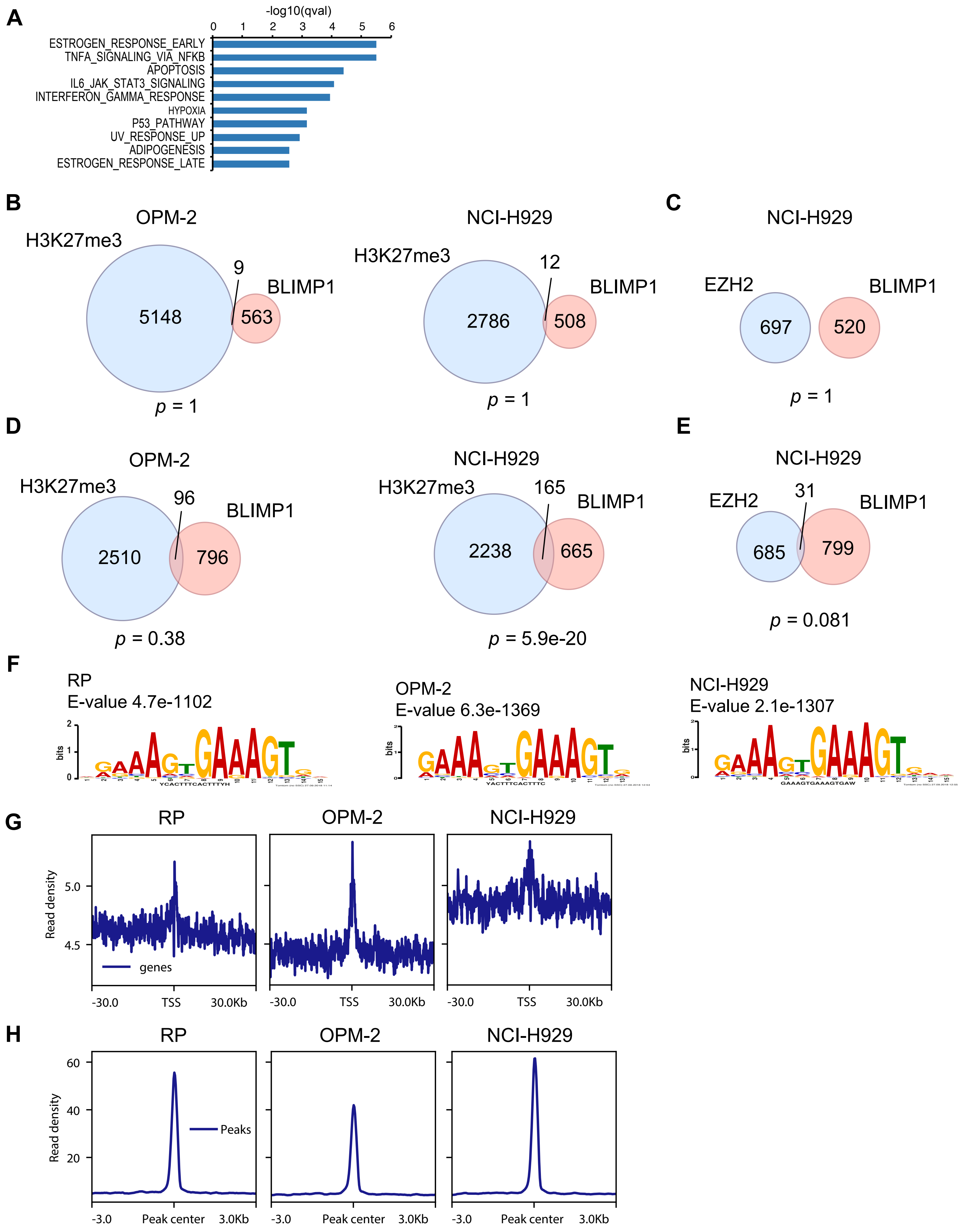


Supplementary Figure 4

(A) Overlap analysis using the Hallmarks gene sets from the molecular signatures database of genes marked by both H3K27me3 and BLIMP1 in the RP cell line, showing the top 10 most significantly enriched gene sets. (B) Venn diagrams of H3K27me3 and BLIMP1 peaks extended ±500 bp, showing overlaps in these regions in the OPM-2 and NCI-H929 cell lines. Not significant as determined by hypergeometric test. (C) Venn diagram of EZH2 and BLIMP1 peaks extended ±500 bp, showing overlaps in these regions in the NCI-H929 cell line. Not significant as determined by hypergeometric test. (D) Venn diagrams of overlapping genes assigned to peaks for H3K27me3 or BLIMP1 in the OPM-2 and NCI-H929 cell lines. Significance determined by Fisher's exact test. (E) Venn diagram of overlapping genes assigned to peaks for EZH2 or BLIMP1 in the NCI-H929 cell lines. Significance determined by Fisher's exact test. (F) Motif enrichment analysis showing the top motif for BLIMP1 ChIPseq experiments in the RP, OPM-2 and NCI-H929 cell lines. (G) Profiles showing enrichment over ±30 kb regions from TSSs assigned to BLIMP1 peaks in the RP, OPM-2 and NCI-H929 cell lines. (H) Enrichment of BLIMP1 signal in the RP, OPM-2 and NCI-H929 cell lines over BLIMP1 peaks from the RP cell line.

#### Supplementary Figure 5

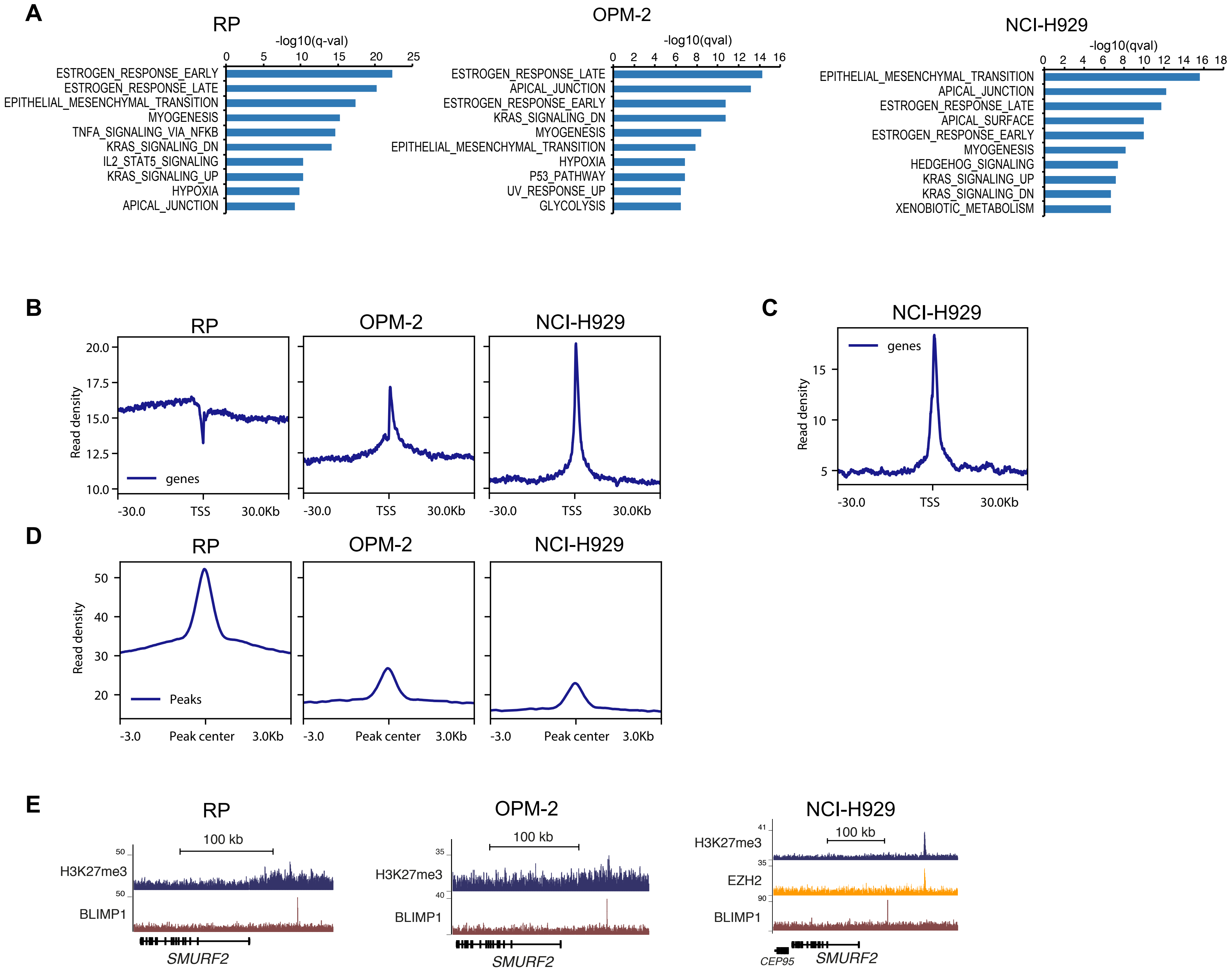


Supplementary Figure 5

(A) Overlaps of H3K27me3 marked genes with Hallmarks gene sets from the molecular signatures database, showing the top 10 most significantly enriched gene sets for the RP, OPM-2 and NCI-H929 cell lines. (B) Profiles showing enrichment over ±30 kb regions from TSSs assigned to H3K27me3 peaks in the RP, OPM-2 and NCI-H929 cell lines. (C) Profile showing enrichment over ±30 kb regions from TSSs assigned to EZH2 peaks in the NCI-H929 cell line. (D) Enrichment of H3K27me3 signal in the RP, OPM-2 and NCI-H929 cell lines over H3K27me3 peaks from the RP cell line. (E) ChIPseq tracks for H3K27me3 and BLIMP1 over the SMURF2 gene in the RP and OPM-2 cell lines, and tracks for H3K27me3, EZH2 and BLIMP1 in the NCI-H929 cell line.

### Supplementary Figure 6

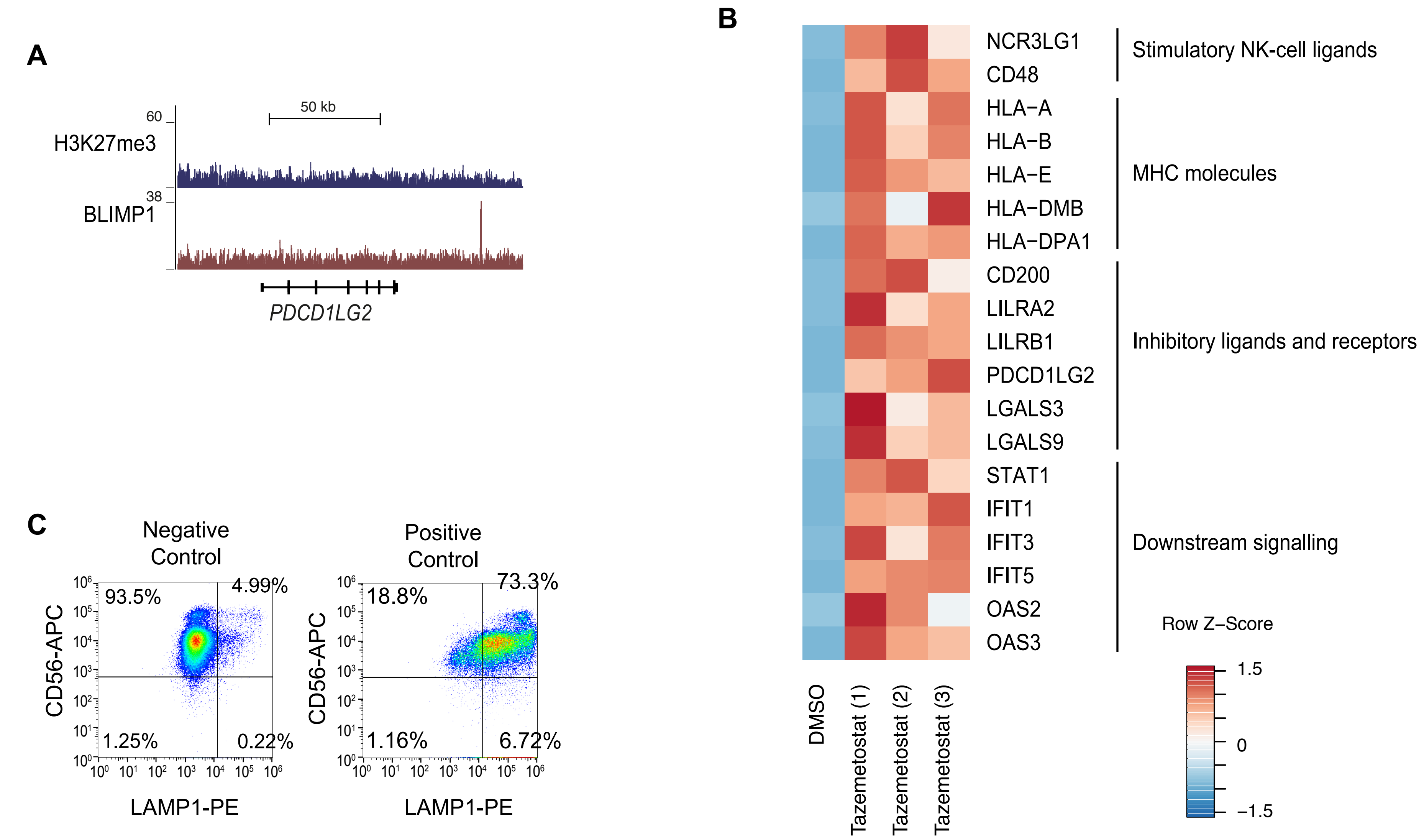


Supplementary Figure 6

(A) ChIPseq tracks for H3K27me3 and BLIMP1 over the PDCD1LG2 gene in the RP cell line. (B) Heat map showing the z-score of the log2 fold change for Tazemetostat compared to DMSO in RP cells with three independent replicates looking at genes involved in stimulation of NK cells, MHC molecules, inhibitory ligands and receptors, and downstream signalling. (C) Negative control for the degranulation assay, consisting of NK cells cultured alone, and positive control consisting of NK cells in co-culture with RP cells treated with 2.5μg/mL phorbol 12-myristate 13-acetate and 0.5μg/mL ionomycin.
