## Supplementary Figure legends and Supplementary materials and methods for "The BLIMP1 – EZH2 nexus in a non-Hodgkin lymphoma"

**Supplementary data**

**Supplementary Data figure legends**

**Supplementary Figure 1**

**(A)** Percentage of live cells as determined by Trypan blue exclusion assay in the RP cells comparing *PmiR1* or *PmiR2* to *NTmiR* following 6 days of induction. **(B)** Annexin V staining of RP cells induced for 48h with dox to express *NTmiR*, *PmiR1* or *PmiR2*; ***p* ≤ 0.01; *** *p* ≤ 0.001 **(C)** Annexin V staining of 48h-induced MW cells with *NTmiR* or *PmiR1*; *p* = 0.0572.

**Supplementary Figure 2**

**(A)** Signal intensity of immunofluorescence staining for BLIMP1 and EZH2 as quantified by CellProfiler from approximately 5000 cells per sample across two independent replicates. **(B)** Relative growth of RP cells treated with tazemetostat for 96h, shown next to RP cells treated with Ibrutinib for 48h.

**Supplementary Figure 3**

**(A)** Graph of RNAseq results depicting normalised transcripts per million (TPM) values for *SMURF2* in RP cells with *NTmiR* and *PmiR1*. Heat maps depicting the Z-score of the log_2_ fold change comparing tazemetostat treatment to DMSO for three independent replicates looking at **(B)** B cell genes and **(C)** apoptosis genes.

**Supplementary Figure 4**

**(A)** Overlap analysis using the Hallmarks gene sets from the molecular signatures database of genes marked by both H3K27me3 and BLIMP1 in the RP cell line, showing the top 10 most significantly enriched gene sets. **(B)** Venn diagrams of H3K27me3 and BLIMP1 peaks extended ±500 bp, showing overlaps in these regions in the OPM-2 and NCI-H929 cell lines. Not significant as determined by hypergeometric test. **(C)** Venn diagram of EZH2 and BLIMP1 peaks extended ±500 bp, showing overlaps in these regions in the NCI-H929 cell line. Not significant as determined by hypergeometric test. **(D)** Venn diagrams of overlapping genes assigned to peaks for H3K27me3 or BLIMP1 in the OPM-2 and NCI-H929 cell lines. Significance determined by Fisher’s exact test. **(E)** Venn diagram of overlapping genes assigned to peaks for EZH2 or BLIMP1 in the NCI-H929 cell lines. Significance determined by Fisher’s exact test. **(F)** Motif enrichment analysis showing the top motif for BLIMP1 ChIPseq experiments in the RP, OPM-2 and NCI-H929 cell lines. **(G)** Profiles showing enrichment over ±30 kb regions from TSSs assigned to BLIMP1 peaks in the RP, OPM-2 and NCI-H929 cell lines. **(H)** Enrichment of BLIMP1 signal in the RP, OPM-2 and NCI-H929 cell lines over BLIMP1 peaks from the RP cell line.

**Supplementary Figure 5**

**(A)** Overlaps of H3K27me3 marked genes with Hallmarks gene sets from the molecular signatures database, showing the top 10 most significantly enriched gene sets for the RP, OPM-2 and NCI-H929 cell lines. **(B)** Profiles showing enrichment over ±30 kb regions from TSSs assigned to H3K27me3 peaks in the RP, OPM-2 and NCI-H929 cell lines. **(C)** Profile showing enrichment over ±30 kb regions from TSSs assigned to EZH2 peaks in the NCI-H929 cell line. **(D)** Enrichment of H3K27me3 signal in the RP, OPM-2 and NCI-H929 cell lines over H3K27me3 peaks from the RP cell line. **(E)** ChIPseq tracks for H3K27me3 and BLIMP1 over the *SMURF2* gene in the RP and OPM-2 cell lines, and tracks for H3K27me3, EZH2 and BLIMP1 in the NCI-H929 cell line.

**Supplementary Figure 6**

**(A)** ChIPseq tracks for H3K27me3 and BLIMP1 over the *PDCD1LG2* gene in the RP cell line. **(B)** Heat map showing the z-score of the _log2_ fold change for Tazemetostat compared to DMSO in RP cells with three independent replicates looking at genes involved in stimulation of NK cells, MHC molecules, inhibitory ligands and receptors, and downstream signalling. **(C)** Negative control for the degranulation assay, consisting of NK cells cultured alone, and positive control consisting of NK cells in co-culture with RP cells treated with 2.5µg/mL phorbol 12-myristate 13-acetate and 0.5µg/mL ionomycin.

**Supplementary materials and methods**

**Suspension cell lines**

RPCI-WM1 (a gift from Prof. Asher A. Chanan-Khan), BCWM.1 (a gift from Prof. Steven P. Treon) and OPM-2 (#ACC50, Leibniz Institute DSMZ) cell lines were maintained in RPMI media (#SH30255.FS, Hyclone, GE Healthcare Biosciences, Denmark) supplemented with 10% foetal bovine serum (FBS) (#SV30160.03, Hyclone, GE Healthcare Biosciences, Denmark). The MWCL-1 (a gift from Stephen M. Ansell) and NCI-H929 (#ACC163, Leibniz Institute DSMZ) cell lines were maintained in RPMI media supplemented with 10% FBS 1mM sodium pyruvate (#SH30239.01, Hyclone GE Healthcare Biosciences) and 50µM beta-mercaptoethanol (#0482, VWR, NY, USA).

**Cloning**

For knock-down of BLIMP1, artificial miRNA sequences were designed to target the *PRDM1* transcript using the RNAiDesigner tool (Thermo Fisher Scientific). These were named *PmiR1* and *PmiR2*. A non-targeting control miRNA (*NTmiR*) was also used as previously described (1). The miRNA sequences were cloned into the tetracycline-inducible pPB-hCMV*1-miR plasmid (1, 2). See Table S20 for miRNA sequences.

Site directed mutagenesis was used to introduce mutations in coding sequence of the FUW-PRD1-BF1 lentiviral expression construct (3) to render it resistant to the targeting miRNAs. The plasmid was amplified using primers listed in Table S21. FUW-EZH2 was derived first by PCR amplification from the pCMVHA-hEZH2 plasmid (4), then incorporated via Gibson assembly (New England Biolabs, Ipswitch, MA, USA) into the FUGW backbone (5). The pCMVHA hEZH2 plasmid was a gift from Kristian Helin (Addgene plasmid # 24230 ; http://n2t.net/addgene:24230 ; RRID:Addgene_24230).

**Construction of knock-down cell lines**

Knock-down cell lines were constructed via electroporation 220V, 350µF using Gene Pulser XCell (Bio-Rad, Hercules, CA, USA) with plasmids containing our miRNAs as described above, using the piggyBac transposon system with the piggyBac transposase, the reverse-tetracycline transactivator and a tetracycline-inducible EGFP, each in a separate plasmid. Stable cell lines were selected for with 0.5 mg/mL or 1 mg/mL G418 disulfate solution (#A6798, Applichem, Darmstadt, Germany) for either the RP and MW, or OPM-2 cells respectively for nine days. The RP cells were seeded at a density of 0.5 cells per well in a 96-well plate and clones with the highest KD were used for further analyses. MW and OPM-2 cells were used as a pool of cells directly after antibiotic selection. miRNA expression was induced with 0.2 or 0.5 µg/mL Doxycyline Hyclate (#sc-211380, Santa Cruz Biotechnology) for either the RP and OPM-2, or MW cells respectively.

**Lentiviral transduction**

Lentivirus was produced by transient transfection of HEK293T cells with the envelope vesicular stomatitis virus G protein (VSVG)-encoding pMD2.G plasmid, the packaging plasmid psPAX2, and either the FUW-PRDI-BF1, FUW-EZH2 or FUGW transfer plasmid. psPAX2 was a gift from Didier Trono (Addgene plasmid #12260; http://n2t.net/addgene:12260; RRID:Addgene_12260). pMD2.G was a gift from Didier Trono (Addgene plasmid #12259; http://n2t.net/addgene:12259; RRID:Addgene_12259).

Viral supernatant was concentrated by ultracentrifugation using 100kDa Amicon centrifugal filters (#UFC910008, Sigma-Aldrich, St. Louis, USA). Suspension cell lines were transduced by spinfection, 800 xg for 30 min with 8 µg/mL polybrene.

**Immunofluorescence staining**

Cells were cytospun onto glass slides for 3 min at 800 rpm. Samples were fixed and permeabilised 10 min in 4% formaldehyde (#28906, Thermo Fisher Scientific), 0.1% triton-X-100 (#T-9284, Sigma-Aldrich), then blocked for 1 h at room temperature with 0.1% triton-X-100, 1% bovine serum albumin (#A1391, Applichem), and 10% goat serum (#16210064, Gibco). Slides were incubated overnight at 4˚C with primary antibodies. Nuclei were counterstained with 1µg/mL DAPI (#sc-3598, Santa Cruz Biotechnology). The secondary antibodies used were goat anti-rabbit Alexa Fluor 647 (#A21244, Thermo Fisher Scientific) and goat anti-mouse Alexa Fluor 546 (#A11030, Thermo Fisher Scientific) at a 1:1000 dilution. The signal intensity within nuclei was quantified using CellProfiler, with background signal subtracted (6).

**Protein extraction**

Cells were lysed in protein sample buffer (60 mM Tris pH 6.8, 2% SDS, 10% glycerol, 0.01% bromophenol blue, 1.25% β-mercaptoethanol), at 95˚C for 5 min. Benzonase nuclease (#sc-202391, Santa Cruz Biotechnology) was used to digest chromatin before loading on SDS-PAGE gel. Histones were extracted using triton extraction buffer (0.5% triton-X-100, 2 mM phenylmethylsulfonyl fluoride, 0.2% NaN_3_ in PBS) and then 0.2 N HCl.

**Antibodies used in western blot and immunofluorescence staining**

The following primary antibodies were used: anti-BLIMP1 (#9115, Cell Signaling Technologies) at a 1:100 dilution for immunofluorescence staining and a 1:1000 dilution for western blot; anti-EZH2 (#612666, BD Biosciences) at a 1:100 dilution for immunofluorescence staining and a 1:1000 dilution for western blot; anti-Actin (#MAB1501, Millipore) at a 1:5000 dilution for western blot; anti-histone H3 (#ab1791, Abcam) at a 1:1000 dilution for western blot; and anti-H3K27me3 (#39536, Active motif) at a 1:1000 dilution for western blot. Proteins were transferred to PVDF membrane (#10600021, GE Healthcare Biosciences) using a wet electrophoretic transfer system (#1703930, Bio-Rad). BLIMP1 was visualised by enhanced chemiluminescence using goat anti-rabbit IgG-HRP secondary antibody (#sc-2004, Santa Cruz Biotechnology) on the ImageQuant LAS 4000 (GE Healthcare Biosciences) following addition of western blotting Luminol reagent (#sc-2048, Santa Cruz Biotechnology). All other proteins were visualised using IRDye800CW goat anti-mouse IgG (#925-32210, LiCor Biosciences GmbH) on the LiCor Odyssey instrument (LiCor Biosciences GmbH). Quantitation of immunoblots was carried out by densitometry using the ImageJ software package (7).

**Resazurin assay**

Resazurin sodium salt (Santa Cruz Biotechnology) was added 5h before the assay end-point, and absorbance measured at 595 and 570nm. Per cent reduction was calculated relative to the fully reduced form of resazurin, resorufin. The percentage reduction of resazurin was calculated according to the following formula:

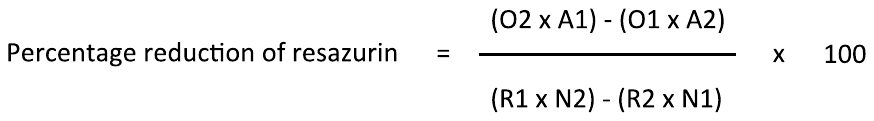

Where O1 = molar extinction coefficient (E) of oxidized resazurin at 570 nm (80586); O2= E of oxidized resazurin at 595 nm  (117216); R1 = E of reduced resazurin (Red) at 570 nm (155677); R2= E of reduced resazurin at 595 nm (14652); A1 = absorbance of test wells at 570 nm; A2 = absorbance of test wells at 595 nm; N1 = absorbance of negative control well (media plus resazurin but no cells) at 570 nm; N2 = absorbance of negative control well (media plus resazurin but no cells) at 595 nm.

**Apoptosis assay**

Apoptosis was assessed using APC-Annexin V (#550474, BD Biosciences) together with Annexin V binding buffer (#556454, BD Biosciences). Staining was carried out according to the manufacturer’s instructions and assessed using the MACSQuant analyser (Miltenyi Biotec).

**cDNA synthesis and RT-qPCR**

cDNA was synthesised using the GoScript Reverse Transcriptase system (#A5001, Promega), in combination with dNTP mix (#R0191, Thermo Fisher Scientific), random primer 6 (#S1230S, New England Biolabs), murine RNase inhibitor (#M0314S, New England Biolabs) and 1.25mM MgCl_2_. RT-qPCR was performed using the SensiFAST SYBR Lo-Rox Kit (#BIO-94020, Bioline) on the 7500 Real-Time PCR system (Applied Biosystems). Relative quantitation was calculated according to the Pfaffl method (8), with *ACTB* and *PPIA* used as reference genes. Primer sequences are listed in Table S22.

**RNAseq**

Total RNA was isolated using TRI-reagent (#AM9738, Thermo Fisher Scientific) according to the manufacturer’s instructions, then treated with HL-dsDNase (#70800-201, ArcticZymes), and purified using the RNeasy MinElute Cleanup Kit (#74204, Qiagen). RNAseq libraries were constructed using the NEB NEBNext® Ultra™ Directional RNA Library Prep Kit for Illumina® (#E7420S, New England Biolabs) in combination with the NEBNext® Poly(A) mRNA Magnetic Isolation Module (#E7490S, New England Biolabs). Pooled samples were clustered on paired-end (PE) flowcells using a cBot instrument (Illumina). Sequencing was performed on Illumina HiSeq 2500 using the v4 SBS sequencing kits with a readlength of 2x125 cycles plus a 6 base index read.  Primary processing and base calling was done using HCS and RTA. Demultiplexing and generation of FASTQ files was performed using scripts from Illumina (bcl2fastq v.1.8).

RNAseq raw reads were trimmed using Trim Galore (<https://www.bioinformatics.babraham.ac.uk/projects/trim_galore/>), and pseudoaligned to the hg38 transcriptome and quantified using Kallisto (9). Differential expression analysis was performed using the likelihood ratio test in Sleuth (10), on a gene level, with a q-value of 0.05, and a log_2_ fold-change of ±0.3 as a significance cutoff. Both condition and batch were used as covariates in the likelihood ratio test. Overlap analysis was performed using the Hallmarks collection of gene sets from the molecular signatures database (11, 12).

**ChIPseq**

ChIP was carried out for BLIMP1, using 60µL rabbit polyclonal antibody recognising the C-terminal region of BLIMP1 (13) with 3×10^7^ nuclei, crosslinked with 0.4% formaldehyde. For EZH2, ChIP was performed using 40µL anti-EZH2 (D2C9, #5246, Cell Signaling Technology) with 2 × 10^7^ nuclei, crosslinked with 1% formaldehyde. Nuclei were sonicated for 6 min for BLIMP1 and 7 minutes for EZH2, using the Epishear™ probe sonicator (#53052, Active Motif) at 25% output with cycles of 15 s on, 30 s off. ChIP was performed for pan-H3 using 3µL anti-histone H3 (D2B12, #4620S, Cell Signaling Technology). For H3K27me3, ChIP was performed using 5µg anti-H3K27me3 (#ab6002, Abcam). All histone ChIP experiments were performed with 1 × 10^7^ cells per ChIP, crosslinked with 0.4% formaldehyde. Histone ChIP samples were sonicated using the Bioruptor® Standard (#UCD-200, Diagenode) 3 x 5 min on high with 30 sec on, 30 sec off. Protein A Dynabeads (#10001D, Invitrogen) were used for all ChIP experiments.

Sequencing was performed as described above. Raw ChIPseq reads were trimmed as above and aligned to the hg38 genome using Bowtie2 with the default settings (14). Peaks were called using MACS2 with q-value cutoffs of 0.01, 0.001 and 0.0001 with the “BAMPE” command for paired-end sequences and otherwise default settings (15), and tracks were visualised using the UCSC genome browser (16). Overlapping regions were identified and tested for significance using the hypergeometric test in the ChIPPeakAnno package in R (17). Overlapping genes were tested for significance using Fisher’s exact test in the GeneOverlap package in R (<http://shenlab-sinai.github.io/shenlab-sinai/>). De novo motif enrichment analysis was performed using MEME ChIP with the input comprising fasta files generated from the regions specified in our peak lists (18). We used the human and mouse HOCOMOCO v11 set of motifs to search for enrichment (19), with default settings. Enrichment of ChIPseq signals over TSSs and peaks was calculated and mapped using Deeptools (20).

**NK cell isolation**

PBMCs were first isolated by density gradient centrifugation with Histopaque-1077 (#10771, Sigma-Aldrich), then NK cells were purified by negative enrichment using the NK cell isolation kit (#130-092-657, Miltenyi Biotech) according to the manufacturer’s instructions. NK cells were cultured overnight in RPMI media supplemented with 10% FBS and 10ng/mL IL-2 (R&D Systems).

**Degranulation Assay**

NK cells were co-cultured with pre-treated RP cells at a 10:1 ratio. Anti-CD107a-PE (clone H4A3, BioLegend) was added to the co-culture media. For the positive control, NK cells in co-culture with RP cells were treated with 2.5µg/mL phorbol 12-myristate 13-acetate and 0.5µg/mL ionomycin. After 1h of co-culture, 2µM monensin was added to every sample and incubated for an additional 4h. Cells were stained with anti-CD56-APC (clone CMSSB, EBioscience), fixed with 1% PFA and analysed on the Sony SH800S flow cytometer. Data were analysed using FlowJo v10.

**Cytotoxicity Assay**

NK cells were co-cultured with pre-treated RP cells at a 20:1 ratio for 4h. Cytotoxicity was assessed by measurement of the adenylate kinase activity in the culture medium using the ToxiLight assay (#LT07-217, Lonza). Luminescence was measured using a Modulus microplate reader (Promega). Values were normalised to wells containing pre-treated RP cells without NK cells, and are presented as relative luminescence units (RLU).

**Supplementary Table S20:** Artificial miRNA sequences

| Name | Sequence (5’-3’) |
| --- | --- |
| *PmiR1* | G**TACACTTCTCTTCAAACTCAG**GTTTTGGCCACTGACTGACCTGAGTTTAGAGAAGTGTACA |
| *PmiR2* | G**TTACTCATCACTCCAATAACC**GTTTTGGCCACTGACTGACGGTTATTGGTGATGAGTAACA |
| Non-targeting (*NTmiR*) | **AAATGTACTGCGCGTGGAGAC**GTTTTGGCCACTGACTGACGTCTCCACGCAGTACATTT |

**Supplementary Table S21:** Primers used for mutagenesis

| FUW-P-miR1-mut_Fwd | gaaaaatgcaCATACATTGTGAACGACCAC |
| --- | --- |
| FUW-P-miR1-mut_Rev | ctcgaattccgCCTCTGTCCACAGAGTCA |

**Supplementary Table S22:** Primers used for qPCR

| **Primer** | **Sequence** |
| --- | --- |
| PRDM1_fwd  PRDM1_rev | GGTACACACGGGAGAAAAGC  GAGATTGCTGGTGCTGCTAA |
| EZH2_fwd  EZH2_rev | ACATCCTTTTCATGCAACACC  GCTCCCTCCAAATGCTGGTA |
| STAT1_fwd  STAT1_rev | CTGTGCGTAGCTGCTCCTT  GAGTCAAGCTGCTGAAGTTCG |
| TFEC_fwd  TFEC_rev | ACATGGGGCTTACAAGTGCT  TCAATGAGGTTGTGGTTGTCC |
| POU2F2_fwd  POU2F2_rev | ACTCATGTTGACGGGCAGC  GGTAGCAGGAACTGAGCAGG |
| MICA_fwd  MICA_rev | ACATTCCATGTTTCTGCTGTTGC  GACCTGCAGGCTCACGA |
| LGALS9_fwd  LGALS9_rev | TCAATGGGACCGTTCTCAGC  GAGGGTTGAAGTGGAAGGCA |
| BCL2L11_fwd  BCL2L11_rev | CCTCGGCGCCCTTTCTT  AGGTTGCTTTGCCATTTGGTC |
| CASP4_fwd  CASP4_rev | TGCTGTTTACAAGACCCACG  AGAGCCCATTGTGCTGTCTC |
| OAS2_fwd  OAS2_rev | CAGGAACCCGAACAGTTCCC  AGGACAAGGGTACCATCGGA |
| PIK3CD_fwd  PIK3CD_rev | TGTACGCCGTGATCGAGAAA  CGGTCTTAAGCTGGTCCTTGT |
| RCAN3_fwd  RCAN3_rev | GCGCGAATAGAACTCCACGA  GCGGCAGGAGATAGGACTTG |
| ZFP36L1_fwd  ZFP36L1_rev | GTCTGCCACCATCTTCGACT  TTTCTGTCCAGCAGGCAACC |
| CIITA_fwd  CIITA_rev | CATCCTTGGGGAAGCTGAGG  CTGTGAGCTGCCTTGGGG |
| TNFRSF14_fwd  TNFRSF14_rev | GCAGTGCCAAATGTGTGACC  CCTGGACGATGCAGAAGTGG |
| HAVCR2_fwd  HAVCR2_rev | GGCTCTTATCTTCGGCGCT  GGGAGGTTGGCCAAAGAGAT |
| PPIA_fwd  PPIA_rev | CATCTGCACTGCCAAGACTGA  TGGCCTCCACAATATTCATGC |
| ACTIN_fwd  ACTIN_rev | AGGCACCAGGGCGTGAT  GCCCACATAGGAATCCTTCTGAC |
